# Structural basis for competitive binding of productive and degradative co-transcriptional effectors to the nuclear cap-binding complex

**DOI:** 10.1101/2023.07.25.550453

**Authors:** Etienne Dubiez, Erika Pellegrini, Maja Finderup Brask, William Garland, Anne-Emmanuelle Foucher, Karine Huard, Torben Heick Jensen, Stephen Cusack, Jan Kadlec

## Abstract

The nuclear cap-binding complex (CBC) co-ordinates co-transcriptional maturation, transport, or degradation of nascent Pol II transcripts. CBC with its partner ARS2 form mutually exclusive complexes with diverse ‘effectors’ that promote either productive or destructive outcomes. Combining Alphafold predictions with structural and biochemical validation, we show how effectors NCBP3, NELF-E, ARS2, PHAX and ZC3H18 form competing binary complexes with CBC and how PHAX, NCBP3, ZC3H18 and other effectors compete for binding to ARS2. In ternary CBCA complexes with either PHAX, NCBP3 or ZC3H18, ARS2 is responsible for the initial effector recruitment but inhibits their direct binding to the CBC. We show that *in vivo* ZC3H18 binding to both CBC and ARS2 is required for nuclear RNA degradation. We propose that recruitment of PHAX to CBC-ARS2 can lead, with appropriate cues, to competitive displacement of ARS2 and ZC3H18 from the CBC, thus promoting a productive rather than a degradative RNA fate.

## Introduction

The heterodimeric nuclear cap-binding complex (CBC, comprising subunits CBP80/NCBP1 and CBP20/NCBP2) binds tightly to the 5′ cap-structure (m^7^GpppNm, where N is the first transcribed nucleotide), a co-transcriptional modification common to all nascent Pol II transcripts. Capping occurs during NELF (negative elongation factor) and DSIF (DRB sensitivity-inducing factor) induced Pol II transcriptional pausing 25-50 nucleotides after initiation (Rambout and Maquat, 2020). The extreme C-terminus of the NELF-E subunit interacts directly with CBC in a groove formed at the CBP20-CBP80 interface and likely plays a role in its recruitment to the capped transcript (Aoi et al., 2020; Narita et al., 2007; Schulze and Cusack, 2017). Upon P-TEFb induced Pol II pause release, during which NELF disassociates, it is likely that ARS2 replaces NELF-E in binding to CBC, through a very similar interaction of its C-terminal region in exactly the same binding groove (Aoi et al., 2020; Schulze and Cusack, 2017). Thus, during early elongation, the CBC-ARS2 (CBCA) complex is formed co-transcriptionally on many, if not all, Pol II transcripts. ARS2 is a multi-domain protein with a largely helical core that includes an RRM domain and projecting N- and C-terminal arms (Melko et al., 2020; O’Sullivan et al., 2015; Schulze and Cusack, 2017). ARS2 has been described to be ‘at the nexus of RNA Pol II transcription, transcript maturation and quality control’ (Lykke-Andersen et al., 2021), and is central to driving transcriptional and post-transcriptional molecular decisions as well as being a ‘general suppressor of pervasive transcription’ (Iasillo et al., 2017; Rouviere et al., 2023). However, how it mechanistically fulfils these functions in a transcript dependent fashion remains incompletely understood.

Many different productive or degradative co-transcriptional ‘classifier’ factors or ‘effectors’, which ultimately contribute to fate determination in a transcript-dependent manner, have been shown to interact competitively and in dynamic exchange with either or both components of CBCA (Andersen et al., 2013; Fan et al., 2017; Giacometti et al., 2017; Hallais et al., 2013; Iasillo et al., 2017; Kiriyama et al., 2009; Muller-McNicoll and Neugebauer, 2014; Schulze and Cusack, 2017) (Garland and Jensen, 2020; Lykke-Andersen et al., 2021; Pabis et al., 2013; Polak et al., 2023; Rambout and Maquat, 2020; Schulze et al., 2018; Winczura et al., 2018). Amongst the productive factors are PHAX (Ohno et al., 2000), NCBP3 (Gebhardt et al., 2015; Schulze et al., 2018), FLASH (Kiriyama et al., 2009; Schulze et al., 2018) and ALYREF (Cheng et al., 2006; Fan et al., 2017; Gromadzka et al., 2016; Viphakone et al., 2019), which promote processing, nuclear transport and export of various RNAs. Primary degradative factors, such as ZC3H18 and ZFC3H1, link CBCA to the exosome-adaptors NEXT (nuclear exosome targeting complex) (Andersen et al., 2013; Lubas et al., 2011; Pabis et al., 2013; Polak et al., 2023) and PAXT (pA tail exosome targeting connection) (Meola et al., 2016) respectively. Both adaptors recruit the ribonucleolytic nuclear exosome, leading to the degradation of aberrant (e.g. 3’ extended snRNAs) or unwanted/excess transcripts (e.g. promoter upstream transcripts (PROMPTs) and enhancer RNAs (eRNAs)) or in some cases to transcript maturation through trimming (e.g. telomerase RNA, some snoRNAs) (Cordiner et al., 2023; Gable et al., 2019; Gockert et al., 2022; Zinder and Lima, 2017). Recently, it has been shown that the transcription restriction factor ZC3H4 can also recruit NEXT to nascent transcripts in an ARS2-dependent fashion (Rouviere et al., 2023). The C-terminal leg domain of ARS2, or its ‘effector domain’, is rich in conserved basic residues and has been implicated in the competitive binding to several effectors, including FLASH, NCBP3, ZC3H18, ZC3H4 and ZFC3H1 (Red1 in *S. pombe*). These effectors all contain a so-called ARS2-recruitment motif (ARM), an acidic-rich short linear motif (SLiM) with the consensus (E/D)(D/E)G(E)(L/I/V) (Dobrev et al., 2021; Foucher et al., 2022; Kiriyama et al., 2009; Melko et al., 2020; Polak et al., 2023; Rouviere et al., 2023; Schulze et al., 2018).

PHAX (phosphorylated adapter for export) is required for intranuclear transport of snRNAs and independently transcribed snoRNAs to Cajal bodies and the CRM1-dependent nuclear export of snRNAs (Boulon et al., 2004; Ohno et al., 2000; Suzuki et al., 2010). PHAX binds directly to the CBC (Ohno et al., 2000; Schulze and Cusack, 2017; Schulze et al., 2018), which requires both CBC subunits (Hallais et al., 2013). A minimal region, PHAX^103-327^, is required for full binding (Schulze et al., 2018), which includes the PHAX non-specific RNA-binding domain (Mourao et al., 2010; Segref et al., 2001). PHAX may initially bind to a wide-range of transcripts (Giacometti et al., 2017) but is thought to be displaced by hnRNP-C on RNAs, such as mRNAs, with long enough (> 200 nts) unstructured 5’ regions (Dantsuji et al., 2023; McCloskey et al., 2012). It has been shown that PHAX binding to CBC is incompatible with the binding of NELF-E (Schulze and Cusack, 2017) and ZC3H18 (Giacometti et al., 2017). However, the precise mode of binding of PHAX to the CBC, and hence how it competes with these factors, is unknown.

NCBP3 was originally described to form an alternative heterodimeric cap-binding complex with CBP80 (Gebhardt et al., 2015), but it was later shown that CBC-NCBP3 form a ternary complex and a quaternary complex with ARS2 as well (Rambout and Maquat, 2021; Schulze et al., 2018). Although NCBP3 does bind the cap-analogue m^7^GTP, presumably via its PARN-like RNA binding-domain, the interaction is significantly weaker than to the canonical CBC (Gebhardt et al., 2015; Schulze et al., 2018). NCBP3 also interacts with the EJC (exon junction complex) and TREX (transcription-export complex), thus positively influencing mRNA export (Dou et al., 2020; Gebhardt et al., 2015). More specifically, NCBP3 has been implicated in the preferential export and translation of certain mRNAs upregulated under stresses, such as viral infection (Gebhardt et al., 2019), hypoxia (Shen et al., 2020) or trimethyl-capped HIV mRNAs (Singh et al., 2022). NCBP3 binding to CBC has been shown to be mutually exclusive with PHAX binding (Schulze et al., 2018).

ZC3H18 was first identified as the factor linking CBCA to the NEXT complex (Andersen et al., 2013) but the protein may have additional functions (Winczura et al., 2018). ZFC3H1 plays a similar role, at least for some PAXT substrates (Meola et al., 2016; Polak et al., 2023), the counterparts in *S. pombe* being Red1 and the MTREC complex (Dobrev et al., 2021; Foucher et al., 2022; Zhou et al., 2015). ZC3H18 binds to the CBCA complex (Winczura et al., 2018) competitively with PHAX (Giacometti et al., 2017), but the structural details underlying this competition are currently unknown.

Here we use insights gained from Alphafold (Jumper et al., 2021) and perform structural, biochemical and biophysical studies to understand how NCBP3, PHAX and ZC3H18 bind to CBCA and how formation of mutually exclusive higher-order CBC complexes occurs. We reveal that the structural basis for competitive binding to the CBC by NCBP3, PHAX and ZC3H18, as well as NELF-E and ARS2, depends on conserved binding modes of these proteins to the same sites on the CBC. We also demonstrate that PHAX, NCBP3 and ZC3H18 compete for binding to the effector domain of ARS2 via their respective ARMs. Indeed, we find that ARS2 is required for initial recruitment of these effectors to CBCA, while preventing their direct binding to the CBC. Finally, for ZC3H18, we show by mutational analysis that its binding to both CBC and ARS2 is required for RNA degradation *in vivo*. Our results suggest that RNA fate determination depends on competition between RNA effectors for sites on both CBC and ARS2, with the outcome being determined by additional transcript-dependent cues.

## Results

### NCBP3 binds to the CBC via its C-terminal tryptophan-containing helix

NCBP3, an RNA-binding domain containing protein of 620 residues (Figure 1A), was previously shown to bind directly to both CBP80 and the complete CBC *in vitro* (Schulze et al., 2018). To identify the CBC-binding region of NCBP3, we used MBP pull-down assays and found that the C-terminal region of NCBP3 (NCBP3^360-620^) retained CBC-binding capacity, while the N-terminal region, NCBP3^1-320^, did not (Figures S1A-S1C). Further truncations showed that NCBP3^560-620^ is sufficient for stable complex formation with CBC (Figures S1B-S1E) and isothermal calorimetry (ITC) measurements determined the *K*_d_ of the CBC-NCBP3^560-620^ interaction to be ∼173 nM, which slightly decreased to ∼115 nM when the CBC was bound to the cap analogue (m^7^GTP) (Figures S1F and S1G).

**Figure 1.**
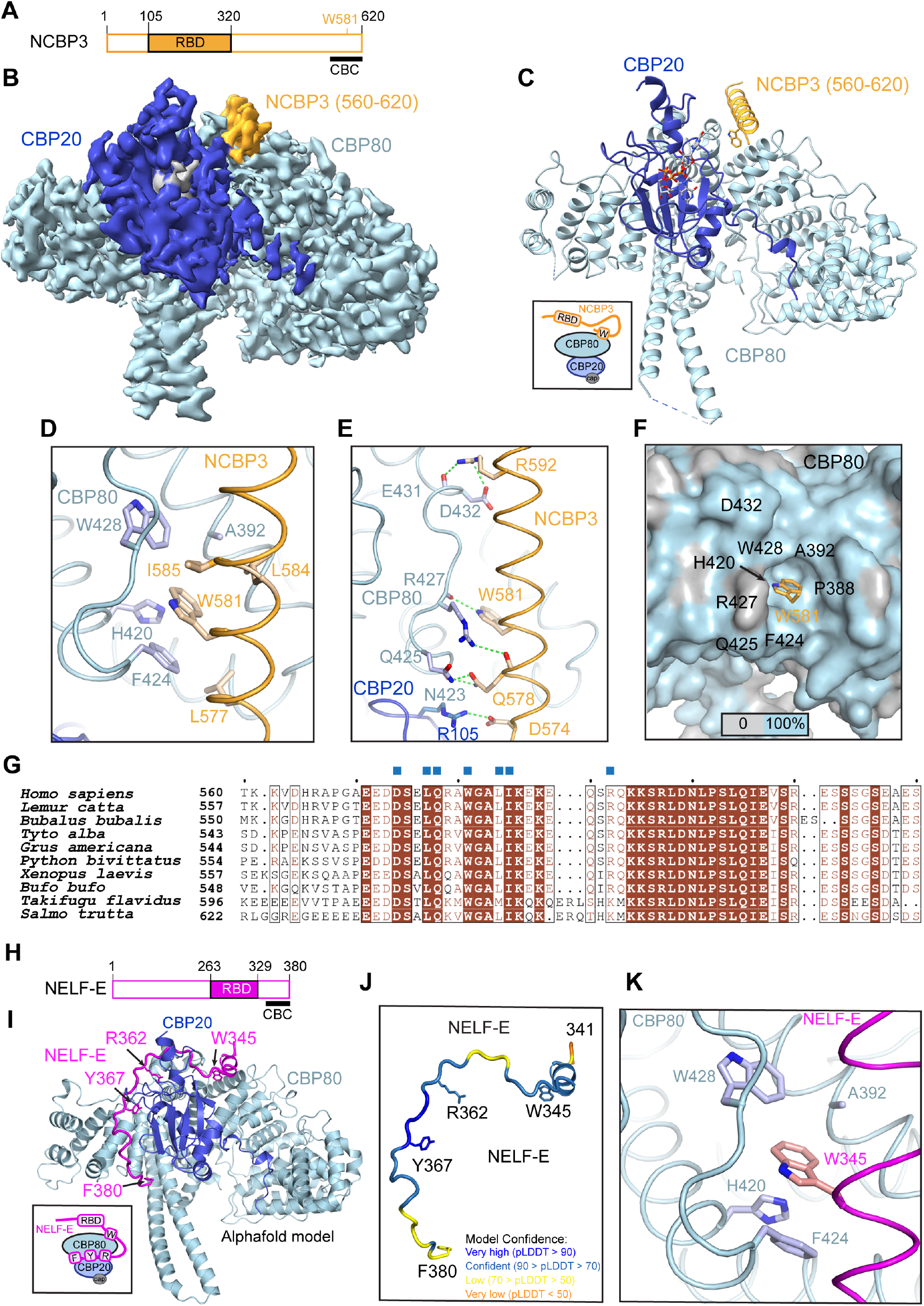
CBC interacts with the tryptophan-containing helix of NCBP3 and NELF-E. **(A)** Schematic representation of the NCBP3 domain structure. RBD: RNA binding domain. The positions of the CBC binding region as identified in this study and W581 are shown. **(B)** Cryo-EM map used to build the CBC-m7GpppG-NCBP3^560-620^ structure. Map is coloured according to molecules location in the structure: CBP80 is in light blue, CBP20 in blue, m^7^GpppG in grey and NCBP3^560-620^ in gold. **(C)** Ribbon representation of the CBC-m^7^GpppG-NCBP3^560-620^ structure. CBP80 is in light blue, CBP20 in blue, m^7^GpppG in grey and NCBP3^560-620^ in gold yellow. W581 is shown as sticks. A schematic model for the interaction of the CBC with the C-terminal Trp-containing helix (W) of NCBP3 is shown in the back square. **(D)** Details of the interaction between the NCBP3 helix (residues 574-594, gold yellow) and CBP80 (light blue). Hydrophobic contacts between NCBP3 and CBP80 residues, including W581, are shown. **(E)** NCBP3 residues form several side and main chain hydrogen bonds and electrostatic interactions with CBP80 and CBP20. **(F)** Surface representation of CBC showing conserved CBP80 residues involved in forming a pocket accommodating NCBP3 W581 (gold yellow). 100 % sequence conservation of surface residues is represented in light blue and is based on CBP80 sequence alignment show in Figure S3D. **(G)** Sequence alignment of NCBP3 proteins covering the C-terminal fragment (residues 560-620) involved in the interaction with CBC. Identical residues are in brown boxes. Blue squares indicate residues involved in the interaction with CBC. **(H)** Schematic representation of the NELF-E domain structure. RBD: RNA binding domain. The CBC binding region as identified in this study is shown. **(I)** Alphafold-multimer predicted CBC-NELF-E complex structure, coloured by chain with NELF-E in magenta. The model was predicted for CBP80, CBP20 and NELF-E^244-380^. Only residues 341-380 of NELF-E are shown. Residues 1-21 of CBP80 predicted with low confidence are not shown. W345, R362, Y367 and F380 of NELF-E are shown as sticks. A schematic model for the interaction of the CBC with the Trp-containing helix (W) and C-terminus containing the conserved arginine (R), tyrosine (Y) and phenylalanine (F) residues of NELF-E is shown in the back square. **(J)** Predicted structure of NELF-E residues 341-380 when bound to CBC, coloured according to pLDDT. **(K)** Details of the interaction between NELF-E W345E and CBP80.

To understand the molecular details underlying this interaction, we determined the structure of the CBC-m^7^GpppG-NCBP3^560-620^ complex by single-particle cryogenic electron microscopy (cryo-EM) at 3.2 Å resolution (Figures 1B and S2, Table S1). This revealed that residues 574-594 of NCBP3 form an α-helix that packs against the second MIF4G domain of CBP80 (Figures 1C and S3A). NCBP3 L577, W581, L584 and I585 bind to a hydrophobic surface on CBP80, with W581 inserting into a pocket comprising CBP80 residues A392, H420, F424 and W428. (Figure 1D). In addition, D574, Q578, W581 and R592 of NCBP3 form main and side chain hydrogen bonds with N423, Q425, R427, E431 and D432 of CBP80 and R105 of CBP20 (Figure 1E). No density was observed for residues 595-620. In agreement with this, removal of C-terminal residues 598-620 resulted in only a slight reduction of the binding affinity (*K*_d_ = 222 nM, Figure S3B). Further truncation to residue 590 increased the *K*_d_ to 6 µM (Figure S3C), suggesting that, in addition to the W581-binding pocket, electrostatic interactions with the conserved positively charged residues 591-597 contribute to the binding. The majority of the CBC and NCBP3 residues involved in these contacts are well conserved across species supporting the importance of this interaction (Figures 1F, 1G and S3D). We note that two very similar isoforms of NCBP3 are reported in many species (including human), which however diverge significantly only beyond the observed CBC-binding site (i.e. beyond QKKSR-597) (Figure S3E). This suggests that both isoforms should be able to bind similarly to the CBC and supports that residues beyond 597 are unlikely to be important for binding.

To confirm a key role of NCBP3 W581 in CBC binding, we generated an NCBP3^560-620^ construct containing the W581E mutation, which did not significantly deviate from the behaviour of the wild-type protein upon gel filtration (Figures S1D, S1E, S3F and S3G. Importantly, using size exclusion chromatography and ITC, we could show that NCBP3^560-620^ ^(W581E)^ no longer binds to the CBC (Figures S3F-S3H). Together, these results demonstrate that CBP80 possesses an interaction surface including a conserved tryptophan-binding pocket (Figures 1D-F) that, in the case of NCBP3, specifically binds to the C-terminal W581-containing helix.

### A tryptophan-containing helix contributes to CBC binding by NELF-E

Since the CBC is involved in multiple, often mutually exclusive, interactions with RNA biogenesis factors, we hypothesised that the CBP80 surface, interacting with the NCBP3 C-terminus, could interact with other competing proteins. NELF-E, a subunit of the NELF complex (Figure 1H), is known to bind the CBC and NELF-E^244-380^ is sufficient for this interaction (Narita et al., 2007), binding to the cap-bound CBC with *K*_d_ ∼50 nM (Schulze and Cusack, 2017). A crystal structure of NELF-E^360-380^ complexed with the CBC showed how the C-terminal extremity of NELF-E binds in the CBP20-CBP80 interface groove (Schulze and Cusack, 2017). However, the binding affinity of NELF-E^360-380^ to the CBC is only 3.3 µM, indicating that upstream sequences of NELF-E likely contribute to the interaction. Furthermore, it was shown that PHAX and NELF-E only compete for binding to the CBC when NELF-E^244-380^, and not NELF-E^360-380^, is used (Schulze and Cusack, 2017).

Examination of the sequence of NELF-E, in conjunction with its AlphaFold (Jumper et al., 2021) predicted structure (AlphaFold-DB, AF-P18615), identified a putative helix with an exposed W345 as a candidate for additional binding to the CBC (Figure S4A). To gain further insight, we used AlphaFold-multimer (Evans et al., 2022) to predict the structure of the CBC-NELF-E complex. We note that AlphaFold consistently predicts the cap-bound form of the CBC, that is with CBP20 fully structured (Mazza et al., 2002), to which the C-terminal peptides of both ARS2 and NELF-E have been shown to have an enhanced binding (Schulze and Cusack, 2017). Furthermore, we confirmed that AlphaFold predicts with high confidence the observed binding of the NCBP3 W581-containing helix to the CBC (data not shown), as well as the binding of the ARS2 C-terminal peptide in the CBP20-CBP80 interfacial groove (data not shown). In the case of the CBC-NELF-E complex, AlphaFold predicts, with high confidence, that residues 344-380 bind to the CBC (Figures 1I, 1J, S4B and S4C). The prediction of the C-terminal 20 residues (360-380) at the CBP20-CBP80 interface corresponds to the experimentally determined CBC-NELF-E structure (Schulze and Cusack, 2017) (Figure S4D). In addition, a short helix (residues 344-350), containing the well-conserved W345, is predicted to bind in the same CBP80 Trp-binding pocket as observed for NCBP3 W581, but, surprisingly, with the opposite polarity to the NCBP3 Trp-containing helix (Figures 1J, 1K and S4E). To confirm this prediction, we mutated NELF-E W345 to glutamate and found that while WT NELF-E^244-380^ bound to the CBC with a *K*_d_ of 113 nM (Figure S4F), the NELF-E^244-380^ (W345E) mutant displayed significantly reduced binding, with a *K*_d_ of 1.3 µM (Figure S4G). The behaviour of NELF-E^244-380^ was not significantly altered by the W345E mutation as judged by gel filtration analysis (Figures S4H-S4J). These results therefore demonstrate, that in addition to its previously described binding to the groove at the CBP20-CBP80 interface, also used by ARS2, NELF-E interacts with the tryptophan-binding pocket used by NCBP3 (Figures 1D and 1F). Binding of this second region likely explains the previously observed discrepancy, that NELF-E^360-380^ does not account for all the binding affinity of NELF-E^244-380^ to the CBC nor to the observed competition with PHAX for CBC binding (Schulze and Cusack, 2017).

### AlphaFold prediction of the CBC-PHAX complex structure

In addition to ARS2, NELF-E and NCBP3, the CBC also binds to PHAX (Figure 2A), an interaction that is critical for the snRNA export pathway (Ohno et al., 2000). In the presence of cap analogue, full-length PHAX binds CBC with a *K*_d_ of 125 nM and PHAX^103-327^, which includes the PHAX RNA binding domain, was proposed to be the minimal construct maintaining almost full CBC binding efficiency (Schulze and Cusack, 2017). The AlphaFold-DB prediction of isolated PHAX (AF-Q9H814), together with sequence alignments, highlighted a conserved W118 on a predicted helix (117-136), positioned within the minimal CBC binding region, which we hypothesised could bind to the CBC in a similar fashion to NCBP3 and NELF-E (Figure S5A). Applying AlphaFold-multimer to the CBC-PHAX complex, PHAX residues 116-133 were confidently predicted to form a helix, containing W118, that would bind to the Trp-binding surface of CBP80 with the same directionality as that of NELF-E (i.e. opposite to the NCBP3 Trp-containing helix) (Figures S5B-S5E). Secondly, the extreme C-terminus of PHAX (390-DLDIF) was predicted to bind at the distal end of the groove between CBP20 and CBP80 in a fashion equivalent to the extreme C-termini of ARS2 (867-DVDFF) and NELF-E (376-LVDGF) (Figures S5B-S5E). Finally, just downstream of the putative Trp helix, PHAX residues 146-153 were predicted to either interact with CBP20 close to the cap-binding site (Figures S5B-S5E) or, less consistently, to the proximal groove between CBP20 and CBP80, depending on whether full-length PHAX or PHAX^101-370^, respectively, were used in the prediction. In the latter case, R147 and Y152 of PHAX interacted in an identical fashion to that observed for the equivalently spaced (RxxxxY) residues in the C-terminal peptides of ARS2 and NELF-E (Figures S5F-S5I). We note that W118, R147, Y152 and F394 are highly conserved amongst metazoan PHAX (Figure 2G), lending credence to their potential functional importance. We also note that the proposed NES of PHAX (133-ATELGILGM) is in this region (Ohno et al., 2000).

**Figure 2.**
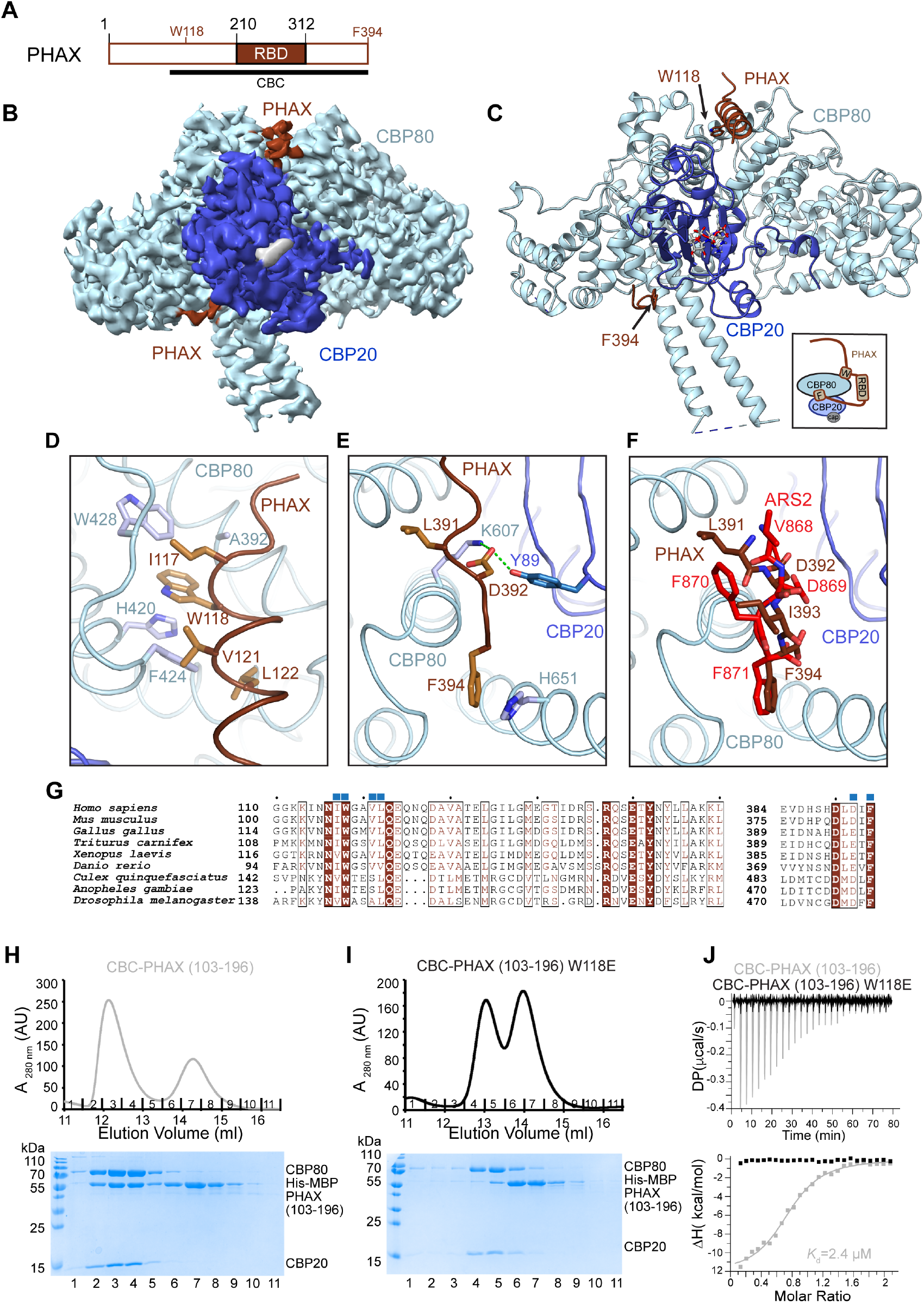
Structural characterisation of the PHAX-CBC complex. **(A)** Schematic representation of the PHAX domain structure. RBD: RNA binding domain. The positions of the CBC binding region as identified in this study, W118 and F394 are shown. **(B)** Cryo-EM map used to build the CBC-capRNA-PHAX structure. Map is coloured according to molecules location in the structure: CBP80 is in light blue, CBP20 in blue, capRNA in grey and PHAX in dark brick. **(C)** Ribbon representation of the CBC-capRNA-PHAX. CBP80 is in light blue, CBP20 in blue, capRNA in grey and PHAX in dark brick. W118 and F394 of PHAX are shown as sticks. A schematic model for the interaction of the CBC with the Trp-containing helix (W) and C-terminus containing the conserved phenylalanine (F) residue of PHAX is shown in the back square. **(D)** Details of the interaction between the PHAX Trp-containing helix (residues 116-133, dark brick) and CBP80 (light blue). Hydrophobic contacts between PHAX and CBP80 residues including W118 are shown. **(E)** Details of the interaction between PHAX C-terminal residues including F394 (dark brick) and the groove between CBP80 (light blue) and CBP20 (blue). **(F)** Comparison of the positioning of the C-termini of PHAX and ARS2 in the groove of the between CBP80 and CBP20. PHAX is in dark brick and ARS2 in red. **(G)** Sequence alignment of PHAX proteins covering region involved in the interaction with CBC. Identical residues are in brown boxes. Blue squares indicate residues involved in the interaction with CBC. **(H)** Superdex 200 gel filtration elution profile and SDS-PAGE analysis of fraction 1-11 of His-MBP-PHAX^103-196^ mixed with CBC, showing co-elution of the three proteins in the first peak. **(I)** Superdex 200 gel filtration elution profile and SDS-PAGE analysis of fraction 1-11 of His-MBP-PHAX^103-196^ (W118E) mixed with CBC, showing lack of binding. **(J)** ITC measurements of the interaction affinity between CBC and WT (grey) or W118E mutant (black) of His-MBP-PHAX^103-196^ in presence of cap analogue.

These predictions suggest that the PHAX and NELF-E binding sites on the CBC overlap in both the Trp-binding pocket and in the CBP20-CBP80 groove. This is consistent with the reported competition between PHAX and NELF-E for binding the CBC (Schulze and Cusack, 2017), and in line with the fact that NELF-E^354-380^ (which does not include W345) does not compete with PHAX. More unexpectedly, the AlphaFold predictions suggested that the PHAX and ARS2 binding sites could also overlap in the CBP20-CBP80 groove. This would imply a potential competition between PHAX and ARS2 binding to the CBC, even though the three proteins are reported to form a ternary complex, denoted CBCAP (Hallais et al., 2013; Schulze and Cusack, 2017). We therefore set out to test these predictions structurally and biochemically.

### Cryo-EM determination of the CBC-PHAX complex structure

We determined the cryo-EM structure of full-length PHAX complexed to the CBC at 3.3 Å resolution (Figures 2B, 2C, S6 and Table S1). We included a 12-mer capped RNA (m^7^GpppAAUCUAUAAUAG) that could engage with both the CBC and potentially the PHAX RNA-binding domain, with the aim of maximally stabilising full-length PHAX binding to the CBC (Ohno et al., 2000). Although in 2D classes, there was fuzzy density that is likely attributable to PHAX (Figure S6), in the processed maps, most of PHAX was not visible, nor was the RNA except for its cap. However, two regions of PHAX gave interpretable density. Firstly, the W118-containing helix binds to CBC (Figure S7A), very much as predicted by AlphaFold (Figures 2B, 2C and S5B-S5E), with W118 inserting into the pocket formed by CBP80 residues A392, H420, F424 and W428 (Figure 2D). In addition, and also as predicted, there is density for the extreme C-terminus of PHAX (390-DLDIF) binding at the very distal end of the CBP80-CBP20 groove similar to ARS2 and NELF-E (Figures 2E and 2F). We also observed extra density near CBP20 Y142 and Y149 and between CBP20 Y50 and CBP80 Y461 (the conserved arginine binding site for ARS2 and NELF-E) but the disjointed nature of the density in this region did not permit to place residues. Thus, neither of the two predicted positions of PHAX^146-153^ (notably the position of the 147-RxxxxY motif) (Figures S5E and S5I) could be confirmed.

To validate and further characterise the mode of binding of PHAX to the CBC, we first verified by gel filtration that PHAX^103-196^ forms a stable complex with the CBC (Figure 2H) and that the W118E mutation abolishes this interaction (Figure 2I). Moreover, the W118E mutation did not significantly alter the behaviour of PHAX^103-196^ as judged by gel filtration analysis (Figure S7B). Using ITC, we measured the *K*_d_ for this fragment to be 2.4 µM (Figure 2J), whereas no binding could be detected for PHAX^103-196(W118E)^ (Figure 2J). The W118E mutation introduced into full length PHAX was not sufficient to abolish binding to CBC in gel filtration (Figures S7C and S7D), consistent with the observed binding to CBC of the PHAX C-terminus (390-DLDIF) and likely also the RNA binding domain (Schulze and Cusack, 2017).

### PHAX, NCBP3 and ZC3H18 contain ARS2 recruitment motifs (ARMs)

Previous studies have shown that ARS2 enhances the binding of PHAX to the CBC and that this might be through a direct interaction of PHAX with ARS2 (Hallais et al., 2013). AlphaFold predicts with high confidence that PHAX 9-EDGQL should bind to the effector domain of ARS2 (Figures 3A, 3B, and S8A-C), exactly as observed crystallographically for the ARS2 recruitment motif (ARM), the ‘EDGEI’ sequence, of *S. pombe* Red1 (Foucher et al., 2022). Sequence alignments of PHAX from metazoans (mammals, birds, fish, amphibians and insects) showed that this putative ARM in PHAX is highly conserved (Figure 3C). Unusually, certain mammals, including human, have a Q in the fourth position, but overall the motif is within the consensus (Foucher et al., 2022). Plant PHAX proteins also contain a putative ARM, either near the N-terminus (EEGEI/L/F, A*. thaliana*-like) or near the C-terminus (EEGEI, maize-like). Consistently, *A. thaliana* PHAX was used in a Y2H assay that demonstrated an interaction between PHAX and ARS2 (Giacometti et al., 2017).

**Figure 3.**
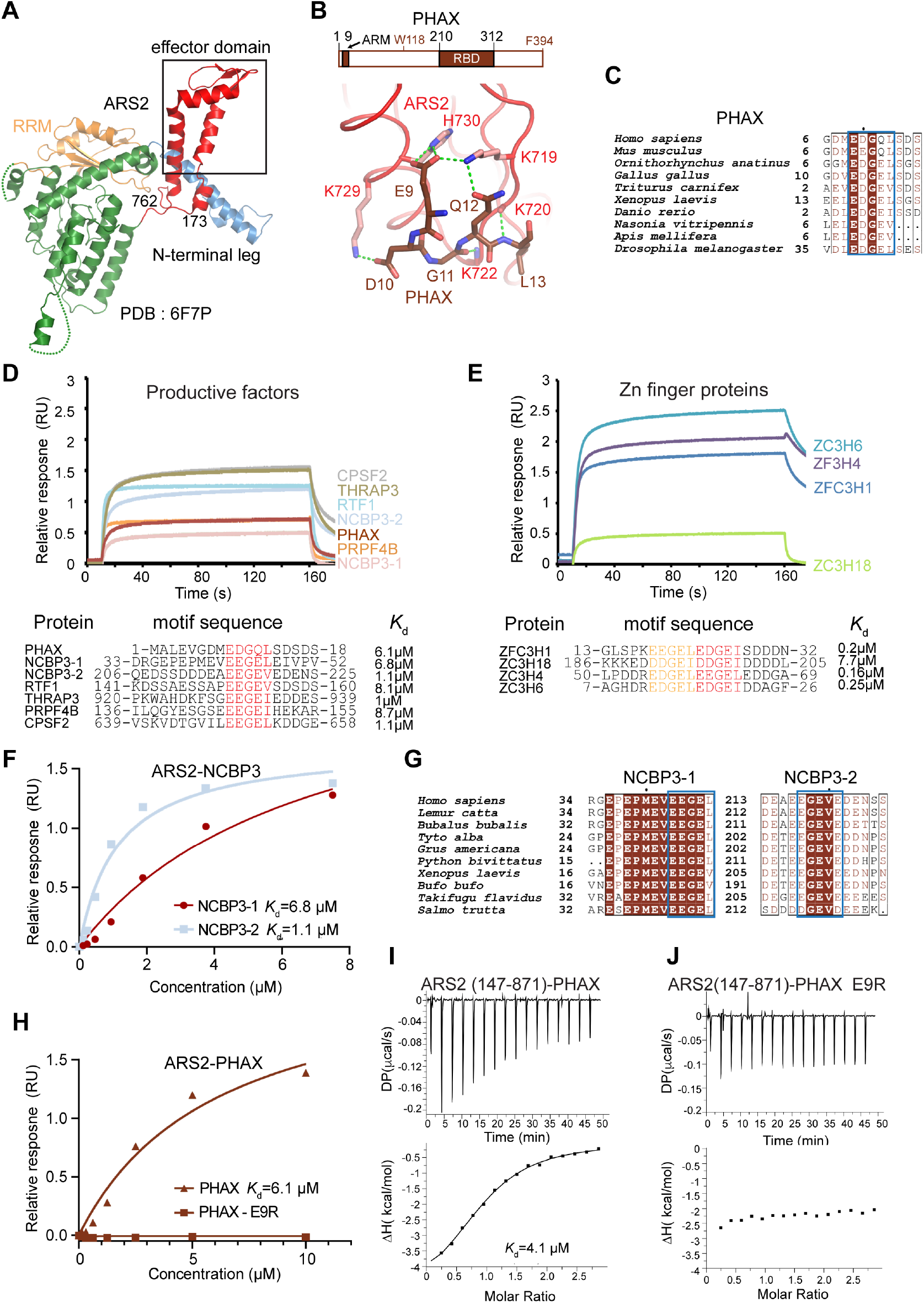
ARS2 interaction with RNA factors. **(A)** Crystal structure of ARS2^173-762^ coloured according to domains (PDB code: 6F7P). The black rectangle highlights the position on the effector domain that interacts with the ARM found in multiple classifier factors. **(B)** Schematic representation of the PHAX domain structure. ARM: ARS2-recruitment motif. Lower panel shows details of the interaction between the ARS2 C-terminal leg and the EDGQ motif of PHAX (residues 9-13, in dark brick) as modelled by AlphaFold-multimer. The model was predicted for full-length ARS2 and PHAX^1-40^. The model with full-length PHAX revealed the same contacts. Corresponding model coloured according to pLDDT and predicted aligned error (PAE) plot are shown in Figures S8A-S8C. **(C)** Sequence alignment of PHAX proteins covering the motif involved in the interaction with ARS2. Identical residues are in brown boxes. The blue rectangle shows the ARM. **(D)** BLI characterization of the interaction between His-ARS2^147-871^ and peptides of RNA-productive factors containing the ARM motif. Representative BLI sensorgrams measured at 1.9-2.5µM concentrations of indicated protein peptides are shown. Sequences of the peptides used as well as the dissociation constants (*K*_d_) are listed the lower panel. **(E)** BLI characterization of the interaction between His-ARS2^147-871^ and peptides of RNA Zn-finger factors containing the ARM motif. Representative BLI sensorgrams measured at the 1.9-2.5µM concentrations of indicated protein peptides are shown. Sequences of the peptides used as well as the dissociation constants (*K*_d_) are listed the lower panel. **(F)** BLI measurement of the binding affinity between His-ARS2^147-871^ and the two NCBP3 ARM motif containing peptides. The *K*_d_ was calculated using the integrated Octet RED96e (ForteBio) software with a 1:1 kinetic model and drawn using Graph Pad Prism5. Measurements were performed in duplicate for each peptide. **(G)** Sequence alignment of NCBP3 proteins covering the two ARM motifs. Identical residues are in brown boxes. The blue rectangles show the ARMs. **(H)** BLI measurement of the binding affinity between His-ARS2^147-871^ and the PHAX WT and mutated ARM motif peptides. The *K*_d_ was calculated using the integrated Octet RED96e (ForteBio) software with a 1:1 kinetic model and drawn using Graph Pad Prism5. Measurements were performed in duplicate for each peptide. **(I)** ITC measurement of the interaction affinity between ARS2^147-871^ and PHAX. **(J)** ITC measurement of the interaction affinity between ARS2^147-871^ and PHAX (E9R).

We measured by Bio-layer interferometry (BLI), the *K*_d_ of ARS2^147-871^ binding to a series of 20-mer peptides, containing the ARMs from a number of human RNA regulatory factors (Foucher et al., 2022), including the PHAX 9-EDGQL motif (within the PHAX^1-18^ peptide), two distinct putative motifs in NCBP3 and double motifs in the Zn-finger containing degradative factors ZFC3H1, ZC3H18 and ZC3H4 (Figures 3D and 3E). *K*_d_s ranged from 0.16 to 9 µM, with the motif context clearly playing a role (Figures 3D, 3E, and S8D-K). Tandem motifs of the tested Zn-finger proteins tended to have higher affinity, perhaps due to avidity affects. The exception was the double motif of ZC3H18 (*K*_d_ of 7.7 µM). However, ZC3H18 in fact contains a third ARM (191-DDGEIDDGEIDDDDLEEGEV), not included in the tested peptide. When binding of all three motifs of ZC3H18 to ARS2^147-871^ was analysed by ITC using MBP-ZC3H18^177-215^, a *K*_d_ of 3.5 µM was measured (Figure S8L). The *K*_d_ decreased to 0.7 µM when the construct MBP-ZC3H18^117-258^, encompassing the downstream Zn-knuckle (residues 222-243), was used (Figure S8M), consistent with the AlphaFold prediction that this region can also interact with the ARS2 C-terminal leg (see below, Figures S9A and S9D). Interestingly, in MBP pull-down assays MBP-ZC3H18^220-353^, still interacted with ARS2^147-871^ even in absence of the ARM motifs providing further support for importance of the Zn-knuckle region in ARS2 binding (Figure S8N, lane 13).

Both putative ARMs from NCBP3 bind ARS2, the second one with higher affinity, and the corresponding motifs are well conserved in vertebrates (Figures 3F and 3G), consistent with a previous study where ARS2 was shown to bind to NCBP3^1-282^ (Schulze et al., 2018). The PHAX motif binds relatively weakly to ARS2 with a *K*_d_ of 6.1 µM, perhaps because of the Q substitution in the fourth position (Figure 3H). We further validated this interaction by first showing that an E9R mutation introduced into the PHAX^1-18^ peptide completely abolished the binding (Figure 3H). Secondly, we confirmed by ITC that full-length PHAX binds to ARS2^147-871^ with a similar *K*_d_ of 4.1 µM (Figure 3I). The E9R mutation essentially abolished the binding of full-length PHAX to ARS2^147-871^ (Figure 3J), indicating that ARM binding to the ARS2 effector domain is crucial for the interaction between the two proteins. The E9R mutation introduced into PHAX did not significantly alter its behaviour as judged by gel filtration analysis (Figures S8O and S8P). These results thus further confirm that the effector domain of ARS2 makes multiple mutually exclusive interactions with productive and degradative factors containing variants of the ARM and demonstrate that such a motif also mediates the interaction between CBC and PHAX.

### ZC3H18 forms a binary complex with CBC as well as with ARS2

ZC3H18 bridges the CBCA to the NEXT complex, since it binds to both CBCA and NEXT components (Andersen et al., 2013; Polak et al., 2023; Puno and Lima, 2022; Winczura et al., 2018). Residues 201-480 of ZC3H18 were identified as containing the major binding site for CBCA, whereas NEXT binds to the C-terminal region of the protein (Figure 4A) (Winczura et al., 2018). As described above, ZC3H18 has three tandem ARMs that we demonstrated biophysically (Figure 3E and Figure S8L) and biochemically (Figure S8N) to bind to ARS2 (Polak et al., 2023). To investigate whether and how ZC3H18 might bind directly to the CBC, we first performed AlphaFold-multimer predictions with the CBC and different fragments of ZC3H18 and the C-terminal region of ARS2. Results are shown for ZC3H18^181-360^ and ARS2^678-871^, which includes both the effector domain and the CBC binding C-terminal peptide (Figure S9). In addition to the expected binding of the C-terminal peptide of ARS2 in the CBP20-CBP80 groove, the predictions consistently and confidently showed: (a) one or the other of the three ZC3H18 ARMs binding in canonical fashion to the top-surface of the effector domain (Figure S9A and S9E), and (b) the ZC3H18 peptide 248-KGNYSL, which is just after the putative Zn-knuckle (residues 222-243), interacting with the shaft of the ARS2 effector domain. These results are consistent with the *in vitro* results described above (Figures S8M, S8N, S9A and S9D).

**Figure 4.**
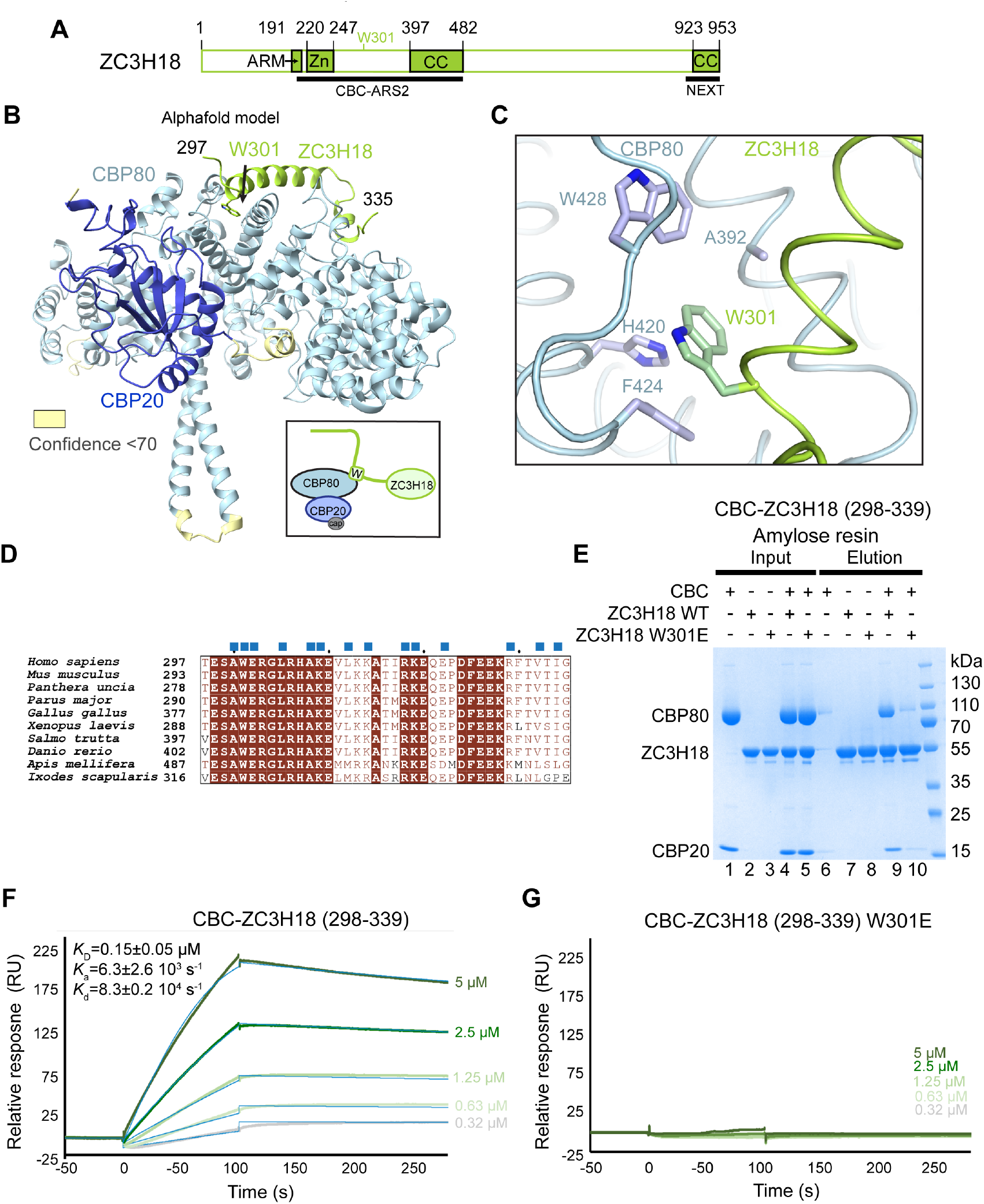
Characterisation of the ZC3H18-CBC complex. **(A)** Schematic representation of the ZC3H18 domain structure. ARM: ARS2-recruitment motif, Zn: Zn-finger, CC-coiled coil. The positions of the previously identified CBC-ARS2 binding region and W301 are shown. **(B)** AlphaFold-multimer predicted structure of the CBC-ZC3H18 complex, coloured by chain with ZC3H18 in lemon. The model was predicted for CBP80, CBP20 and ZC3H18^277-356^. Only residues 297-335 of ZC3H18 are shown. Residues 1-21 of CBP80 predicted with low confidence are not shown. W301 of ZC3H18 is shown as sticks. Residues with pLDDT below 70 are highlighted in yellow. The model coloured according to pLDDT and the corresponding predicted aligned error (PAE) plot are shown in Figures S10B-S10D. A schematic model for the interaction of the CBC with the Trp-containing helix (W) of ZC3H18 is shown in the back square. **(C)** Details of the interaction between the ZC3H18 helix (residues 300-321, lemon) and CBP80 (light blue). Hydrophobic contacts between ZC3H18 W301 and the CBP80 residues are shown. **(D)** Sequence alignment of ZC3H18 proteins covering the region involved in the interaction with CBC (residues 297-335). Identical residues are in brown boxes. Blue squares indicate residues involved in the interaction with CBC according to the AlphaFold model. **(E)** MBP pull-down experiments of WT and W301E mutant of His-MBP-ZC3H18^298-339^ with CBC. All proteins were first purified by Ni^2+^-affinity chromatography. Proteins were mixed as indicated above the lanes. A total of 1% of the input (lanes 1–5) and 10% of the eluates (lanes 6–10) were analysed on 4-20% gradient SDS-PAGE gels stained with Commassie brilliant blue. CBC is retained by ZC3H18^298-339^ (lane 9), whereas its W301E mutation significantly reduces the CBC binding (lane 10). **(F, G)** Binding kinetics of the CBC complex, in 2-fold dilutions, to WT His-MBP-ZC3H18^298-339^ shown in **F** and the W301E mutant of His-MBP-ZC3H18^298-339^ shown in **G**, captured on NTA sensor chip by surface plasmon resonance. Sensorgrams recorded for the association (100 s) and dissociation (200 s) phases for the various CBC concentrations are shown as solid green lines, with the corresponding curves fitting to a 1:1 Langmuir binding superimposed as solid blue lines. The derived kinetic rate constants are shown in **F**.

In addition, consistently and with high confidence, ZC3H18^297-335^ was predicted to bind to CBP80 (Figures 4B, S9A, S9G and S10A-D) with the conserved W301 in the Trp-binding pocket also competed for by NELF-E, PHAX and NCBP3 (Figures 4C and 4D). Trp301 is at the N-terminus of a helix (300-321) that packs against CBP80 and projects in the same direction as the NCBP3 binding helix but at a different angle (Figure 4B). A second helix (residues 325-331) also interacts with CBP80. Predictions with larger fragments did not highlight any other regions of ZC3H18 that might bind to other sites on the CBC, notably the groove-binding site, although this cannot be excluded.

To test this prediction, we performed MBP pull-down experiments with His-MBP-ZC3H18^298-339^. The WT construct was able to pull-down the CBC, whereas binding of the W301E mutant was significantly reduced (Figure 4E, lanes 9,10). We then measured the *K*_d_ between His-MBP-ZC3H18^298-339^ and the CBC using surface plasmon resonance (SPR) and obtained a value of 0.15 µM (Figure 4F), whereas binding of the W301E mutant was not detectable (Figure 4G). The W301E mutation did not significantly alter the behaviour of this ZC3H18^298-339^ fragment as judged by gel filtration analysis (Figures S10F-S10H). Further evidence for this interaction was obtained by a crosslink between ZC3H18 K319 and CBP80 K221 reported in a study on the spliceosome (Townsend et al., 2020), which is perfectly compatible with the AlphaFold prediction (Figure S10E). In summary, these results clearly show that the contacts between ZC3H18 and either the CBC or ARS2 directly overlap with those of the positive RNA biogenesis factors NCBP3 and PHAX.

### In ternary complexes, ARS2 inhibits direct binding of PHAX, NCBP3 and ZC3H18 to CBC

PHAX was previously shown to form a ternary complex with CBC-ARS2 (Hallais et al., 2013; Schulze and Cusack, 2017). Since the binding sites for ARS2 and PHAX within the CBP20-CBP80 groove overlap, suggesting possible competition between ARS2 and PHAX for CBC binding, we further investigated interactions between these three proteins. Using size exclusion chromatography we found, in agreement with our ITC results, that ARS2^147-871^ and PHAX indeed interact and that the ARM of PHAX is required for this interaction, since the E9R PHAX mutant no longer bind ARS2 (Figures 5A-5C). We could further show that the E9R PHAX mutation does not affect the binding of PHAX to the CBC, as that interaction is mediated by at least the W118-containing helix and the PHAX C-terminus (Figures 5D-5F). Importantly, while WT PHAX forms a ternary complex with CBC-ARS2 as expected, the E9R PHAX mutant is unable to interact with CBC-ARS2, despite the presence of its CBC-binding regions (Figures 5G-5I). In addition, when CBC is bound by ARS2, it can no longer interact with NCBP3^560-620^, which lacks the NCBP3 ARM motifs (Figures 5J and 5K).

**Figure 5.**
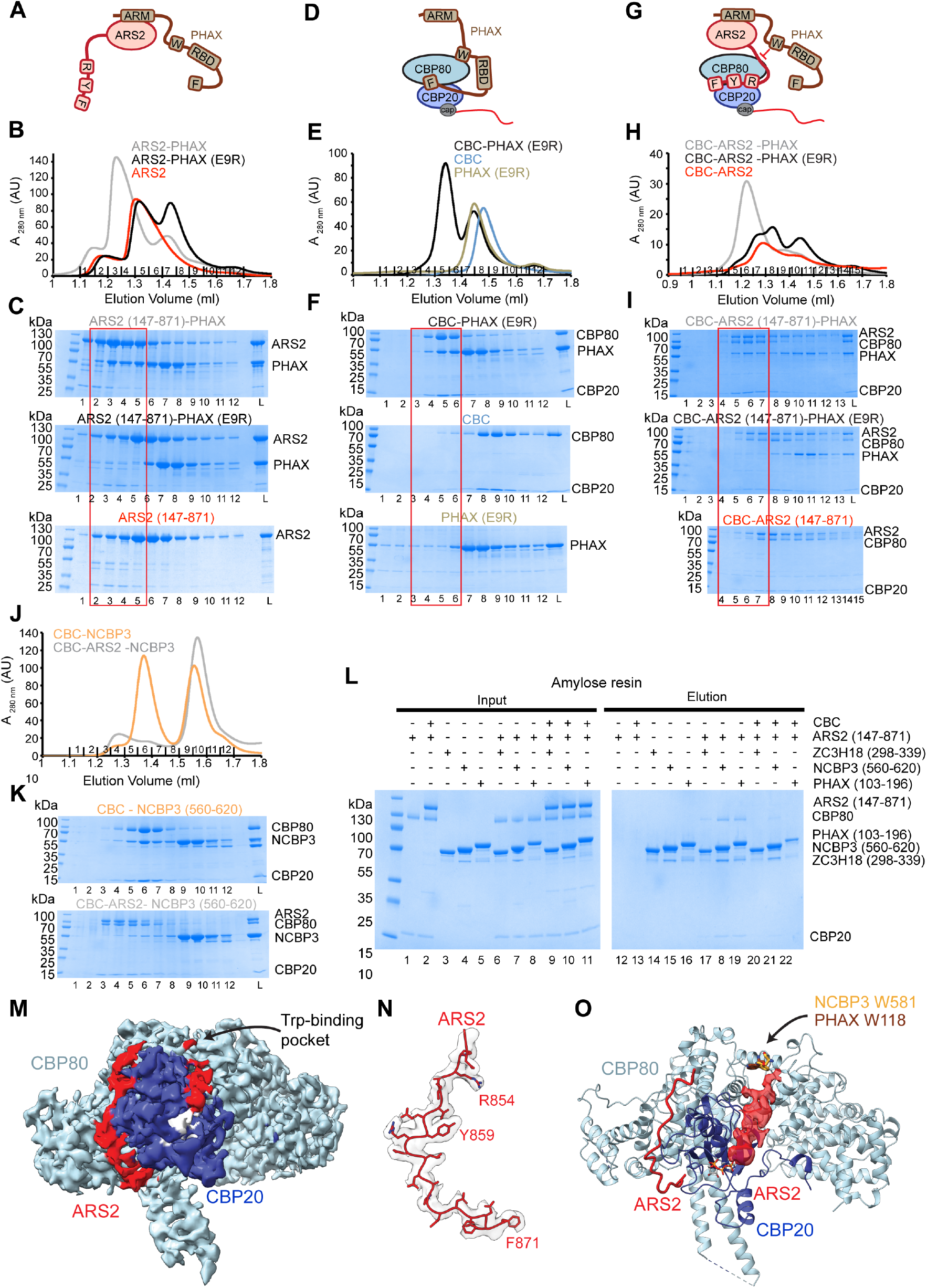
Competition between ARS2 and RNA classifier factors for CBC binding. **(A)** Schematic representation of the ARS2-PHAX interaction. ARS2 residues R852, Y859 and F871 mediating the interaction with CBC are shown as boxes labelled R, Y and F. PHAX ARS2-binding EDGQL motif (ARM), W118-containing helix (W) and F394-containing C-terminus (F) as well its RBD domain are as shown as boxes. PHAX interacts with ARS2 via its N-terminal ARM motif. **(B)** Overlay of Superdex 200 gel filtration elution profiles of ARS2^147-871^ and its individual mixtures with WT PHAX and with the E9R PHAX mutant. While WT PHAX co-elutes with ARS2^147-871^, the E9R PHAX mutant does not. **(C)** SDS-PAGE analysis of fraction 1-12 of the Superdex 200 gel filtration elution profiles shown in **B**. The red square indicates fractions containing the ARS2-PHAX complex compared to elution profiles of ARS2^147-871^ and its mixture with PHAX E9R. **(D)** Schematic representation of the CBC-PHAX interaction. PHAX binds CBC via its W118-containing helix (W) and F394-containing C-terminus (F). **(E)** Overlay of Superdex 200 gel filtration elution profiles of CBC, PHAX E9R and their mixture in presence of m^7^GTP. Despite the E9R mutation, PHAX still binds CBC. **(F)** SDS-PAGE analysis of fraction 1-12 of the Superdex 200 gel filtration elution profiles shown in **E**. The red square indicates fractions containing the CBC-PHAX complex compared to elution profiles of individual proteins. **(G)** Schematic representation of the interaction between CBC-ARS2 and PHAX. PHAX binds ARS2 via its N-terminal ARM motif. ARS2 binding to CBC prevents the direct binding of PHAX to CBC, **(H)** Overlay of Superdex 200 gel filtration elution profiles of CBC-ARS2^147-871^ complex and its mixture with WT PHAX or PHAX E9R of presence of m^7^GTP. While WT PHAX co-elutes with CBC-ARS2^147-871^, the E9R PHAX mutant does not. **(I)** SDS-PAGE analysis of fraction 1-12 of the Superdex 200 gel filtration elution profiles shown in **H**. The red square indicates fractions containing the CBC-ARS2-PHAX complex compared to elution profiles of CBC-ARS2^147-871^ and its mixture with PHAX E9R. **(J)** Overlay of Superdex 200 gel filtration elution profiles of NCBP3^560-620^ mixed with CBC or CBC-ARS2^147-871^ in presence of m^7^GTP. While NCBP3^560-620^ co-elutes with CBC, in presence of ARS2^147-871^ it elutes in a peak separate from CBC-ARS2^147-871^. **(K)** SDS-PAGE analysis of the Superdex 200 gel filtration elution profiles shown in **J.** **(L)** MBP pull-down experiments of His-MBP tagged ZC3H18^298-339^, NCBP3^560-620^ and PHAX^103-196^ with CBC and the CBC-ARS2^147-871^ complex in presence of m^7^GTP. All proteins were first purified by affinity chromatography. Proteins were mixed as indicated above the lanes. A total of 2% of the input (lanes 1–11) and 40% of the eluates (lanes 11–22) were analysed on 4-20% gradient SDS-PAGE gels stained with Commassie brilliant blue. CBC is retained by all ZC3H18^298-339^, NCBP3^560-620^ and PHAX^103-196^ (lane 17-19), but when it is first bound by ARS2^147-871^ its levels are significantly reduced (lanes 20-22). **(M)** Cryo-EM map used to build the CBC-m^7^GTP-ARS2^147-871^ structure. The map is coloured according to each molecule’s location in the structure: CBP80 is in light blue, CBP20 in blue, m^7^GTP in grey and ARS2 in red. **(N)** Cryo-EM density for the ARS2^852-871^ peptide within the CBC-m^7^GTP-ARS2^147-871^ complex. **(O)** Interpretation of density map for ARS2-CBC interaction. Density 2.5 Å around the CBC-ARS2 model was subtracted to only highlight additional density spanning from the CBP20 cap-binding site to the CBP80 Trp-binding site (the surface is shown in red). For comparison, NCBP3 W581 and PHAX W118 are shown in their respective positions (within their complexes with CBC) as sticks.

ARS2 interference with PHAX and NCBP3 binding to the CBC could also be observed in pull-down assays. Here, NCBP3^560-620^ and PHAX^103-196^, that lack their respective ARMs, did interact with the CBC but not with CBC-ARS2^147-871^ (Figure 5L, lanes 20,21). In addition, these experiments revealed that the presence of ARS2 also blocks the binding of ZC3H18^298-339^ to the CBC (Figure 5L, lane 22). Taken together, these results suggest that PHAX, NCBP3 and ZC3H18 are initially recruited to CBCA only via their ARMs while their direct binding to the CBC, in the presence of ARS2, is severely weakened or even blocked, perhaps through steric hindrance of Trp-helix binding to the CBC.

To explore this hypothesis further, we determined the cryo-EM structure of CBC-m^7^GTP-ARS2^147-871^ at 3.4 Å resolution (Figure S11). While the bulk of ARS2 was not resolved, presumably because of its flexible connection to the CBC, we found that the C-terminus of ARS2 binds in the CBP20-CBP80 groove as previously reported (Schulze et al., 2018) (Figures 5M and 5N). However, we also observed additional density extending down from the CBP20 cap-binding site towards the Trp-binding pocket (Figures 5M and 5N). Although the resolution did not permit assignment of this density to particular residues of ARS2 (and Alphafold does not confidently predict this interaction), occupancy of this region by ARS2 would compete with Trp-helix binding by several factors, as observed biochemically.

### The conserved W301 of ZC3H18 is required for stable association with CBC and for function in NEXT-mediated RNA decay

Consistent with the *in vitro* interaction studies described above, we have previously demonstrated that the ARM is required for the stable binding of ZC3H18 to ARS2 *in vivo*, as also observed for the ZFC3H1 and ZC3H4 proteins (Polak et al., 2023; Rouviere et al., 2023). To expand on this, we explored the contribution of the conserved tryptophan site in ZC3H18 in the assembly of CBCA-ZC3H18 complexes with the aim of addressing the functional consequences of disrupting both the ARM and tryptophan sites *in vivo*. To achieve this, we utilized a mouse embryonic stem (mES) cell system where endogenous ZC3H18 alleles were tagged with a mini-auxin-inducible degron and 3xFLAG (3F-mAID) tag (Gockert et al., 2022; Natsume and Kanemaki, 2017; Rouviere et al., 2023), allowing the rapid depletion of ZC3H18-3F-mAID protein upon addition of indole-3-acetic acid (IAA). We then used the Piggybac transposon system (Ding et al., 2005) to stably integrate MYC-tagged ZC3H18-WT, ZC3H18-W297E (mouse equivalent of W301E) or a ZC3H18-ARMmut (D188A, E190A, D193A, E195A, E203A, E205A) (Figure 6A) with equivalent mutations previously shown to disrupt the connection with ARS2 (Polàk et al., 2023). We carried out 16 hr IAA (+) or control (-) treatments, ensuring a complete depletion of ZC3H18-3F-mAID (Figure 6B). To address the functional capacity of the ZC3H18 variants in NEXT-mediated RNA decay, known NEXT targets were analysed by RT-qPCR. Consistent with previous observations, ZC3H18 depletion increased levels of NEXT-sensitive PROMPTs (Figure 6C) (Polak et al., 2023; Rouviere et al., 2023; Winczura et al., 2018), which was rescued by complementation with WT MYC-ZC3H18 (Figure 6C). In contrast, both ZC3H18-W297E and ZC3H18-ARMmut were unable to complement the ZC3H18-depletion phenotype, suggesting both mutants are non-functional (Figure 6C).

**Figure 6.**
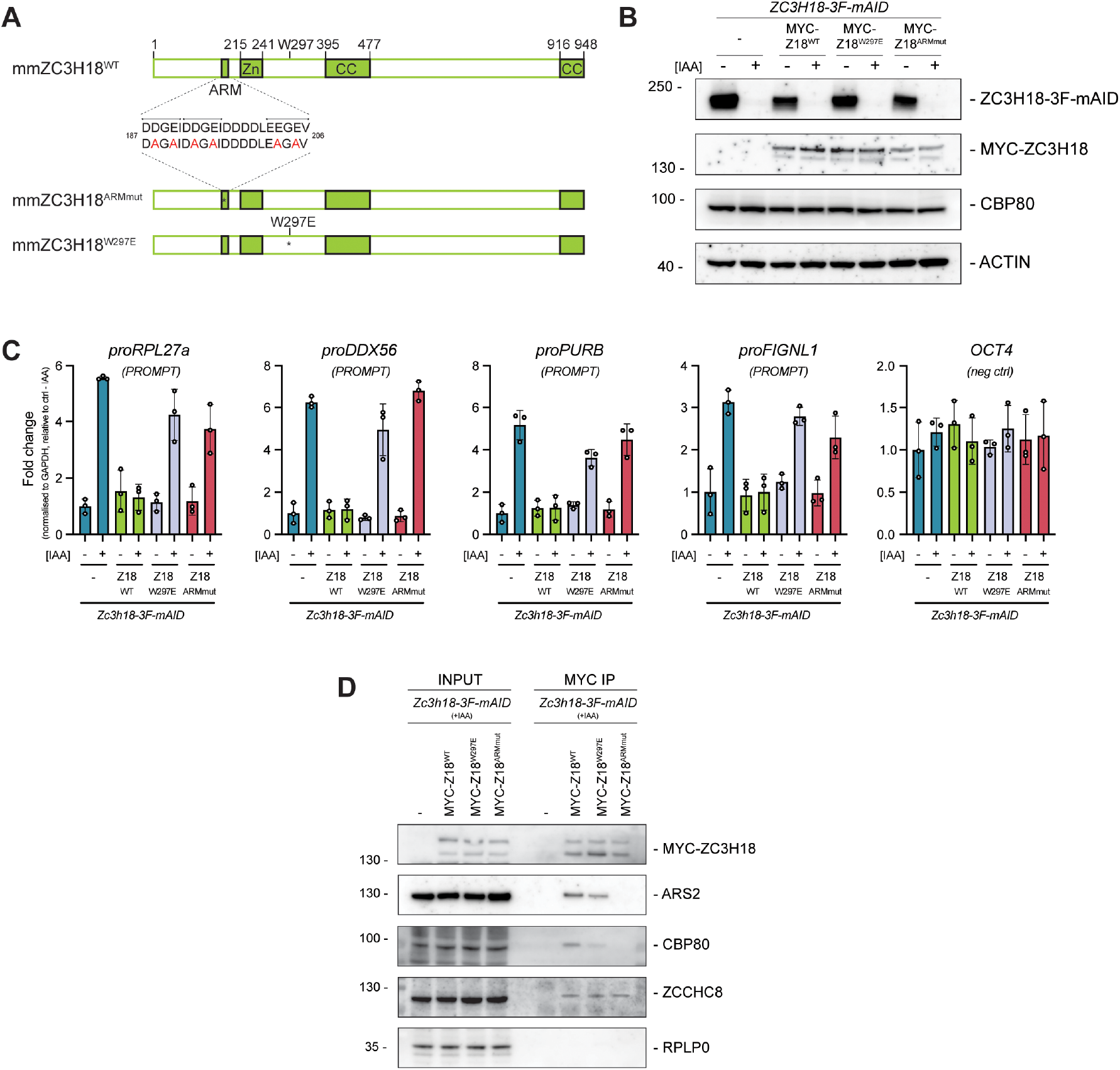
Functional analysis of ZC3H18 ARM and Trp mutants in cells. **(A)** Schematic representation of ZC3H18, depicting its predicted domains, and the mutations used in this study. CC = coiled coil, Zn = zinc finger. The ARS2-recruitment motif (ARM) is shown and zoomed in sequence displays diagnostic point mutations introduced in the ZC3H18^ARMmut^. The position of W297E is indicated with an asterisk (*). **(B)** Western blotting (WB) analysis of lysates from Zc3h18-3F-mAID mouse embryonic stem (mES) cell lines stably expressing the indicated MYC-tagged ZC3H18 mutants. Samples were either mock treated (-) or treated with auxin (IAA) for 16 hours (+) to deplete Zc3h18-3F-mAID protein. Membranes were probed with antibodies against FLAG, MYC, NCBP1 and ACTIN as a loading control. **(C)** RTqPCR analysis of NEXT-sensitive RNAs (proRPL27a, proDDX56, proPURB, proFIGNL1). Total RNA was isolated from *Zc3h18-3F-mAID* mES cell lines stably expressing MYC-ZC3H18^WT^, -ZC3H18^W297E^ or -ZC3H18^ARMmut^ and with the parental cell line (-) as a negative control. Cells were treated as in panel **B**. Results were normalized to GAPDH mRNA levels and plotted relative to Zc3h18-3F-mAID – IAA control samples. Columns represent the average values of biological triplicates per sample with error bars denoting the standard deviation. Individual data values from replicates are indicated as points. **(D)** WB analysis of MYC IPs from lysates of *Zc3h18-3F-mAID* cells stably expressing MYC-tagged ZC3H18^WT^, ZC3H18^W297E^ or ZC3H18^ARMmut^ and with the parental cell line (-) as a negative control. Cells were treated with IAA (16h) to deplete endogenous Zc3h18-3F-mAID protein. Input (left panel) and IP (right panel) samples were probed with antibodies against MYC, ARS2, NCBP1, ZCCHC8 and RPLP0 as a loading control.

To address molecular interactions of the mutants *in vivo*, we performed MYC-immunoprecipitation (IP) experiments from ZC3H18-3F-mAID cells, expressing MYC-ZC3H18-WT, ZC3H18-W297E or ZC3H18ARMmut variants and assessed their interactions with ARS2, CBP80 and ZCCHC8. Samples were depleted of endogenous ZC3H18-3F-mAID for 16 hr to avoid any complications due to the dimeric nature of the NEXT complex (Gerlach et al., 2022:Puno, 2022 #294). MYC-ZC3H18-WT efficiently co-purified ARS2, CBP80 and ZCCHC8 (Figure 6D). Upon mutation of the ZC3H18 ARM, ARS2 and CBP80 interactions were both abrogated, consistent with previous observations in HeLa cells (Polak et al., 2023), whilst, as expected, the ZCCHC8 interaction remained intact. Finally, ZC3H18-W297E co-purified ARS2 but displayed a reduction in CBP80 coIP levels as compared to the MYC-ZC3H18WT control (Figure 6D), suggesting that the W297E mutant is unable to fully engage in stable association with the CBC.

Taken together, these results show that ARS2 plays the dominant role in bridging CBC to ZC3H18 and hence the NEXT complex, but that the direct interaction between CBC and ZC3H18 strengthens the ternary complex and is required for NEXT-mediated degradation.

## Discussion

The relatively large size of the CBC compared with other cap-binding proteins has long been thought to be a consequence of its function as a platform for binding of multiple, competing co-transcriptional factors that contribute to transcript fate determination. However, structural information on how many of these factors interact with the CBC and its cofactor ARS2 has been lacking. In this work, we have used Alphafold predictions and cryo-EM structure determination together with biochemical and biophysical analyses to characterise the binary complexes of effectors NCBP3, PHAX and ZC3H18 with firstly the CBC and secondly ARS2, as well as the ternary complexes of these same factors with CBCA. These results reveal an intriguing situation whereby each of these effectors (and possibly others) compete for the same binding sites on both ARS2 and the CBC, presumably in dynamic flux (Giacometti et al., 2017), until some transcript dependent cue stabilises one complex over another to commit RNA metabolism in a specific direction.

Concerning binary interactions with the CBC, including results presented here, notably the identification of a conserved Trp binding pocket on CBP80, five distinct, non-overlapping interaction sites on the CBC have now been identified (Figures 7A and 7B). These include: (1) the proximal CBP20-CBP80 interfacial groove binding site, where at least NELF-E and ARS2 bind, each with a RxxxxY motif, (2) the distal groove site, where C-terminal peptides ending in phenylalanine (…xDxF-Cter) were shown to bind for ARS2, NELF-E and PHAX; (3) the tryptophan-helix binding site on CBP80, where at least NELF-E, NCBP3, PHAX and ZC3H18 interact, each with a conserved tryptophan bound in the same CBP80 pocket irrespective of the directionality of the helix; (4) the peroxisome proliferator-activated receptor-γ coactivator 1α (PGC-1α) binding site on CBP80 (PDB code: 6D0Y) (Cho et al., 2018; Rambout et al., 2023) and (5) the Transcription elongation regulator 1 (TCERG1) binding site, where, in the human pre-Bact-1 spliceosome structure (PDB code: 7ABG), one of the FF domains binds to CBP80 (Townsend et al., 2020).

**Figure 7.**
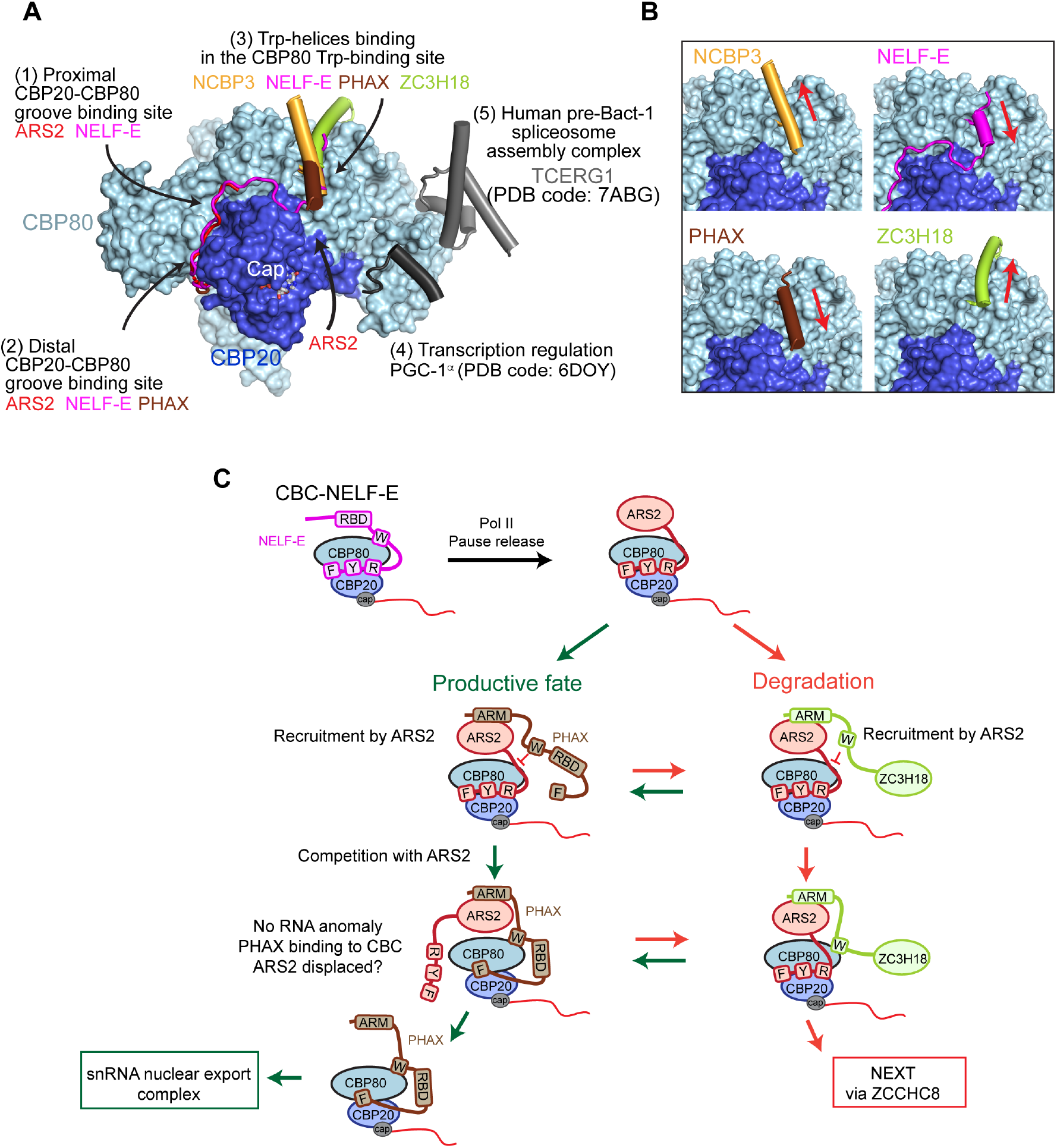
Model for CBC-ARS2 competing interactions. **(A)** Summary of structurally characterised CBC interactions. **(B)** Comparison of CBC interactions with the Trp-containing helices of NCBP3, NELF-E, PHAX and ZC3H18. The red arrows indicate the helix direction. **(C)** Model for the competing interactions of NELF-E, ARS2, PHAX and ZC3H18 with CBC. Following the RNA polymerase II pause release NELF-E is replaced on CBC by ARS2. Both proteins interact with CBC with their C-termini comprising conserved arginine (R), tyrosine (Y) and phenylalanine (F) residues. NELF-E binds CBC also with its Trp-containing helix (W). Productive factors such as PHAX (and NCBP3) as well as degradative factors such as ZC3H18 are recruited to CBC-ARS2 by via their ARM motif binding to the C-terminal leg of ARS2. At this stage, their direct binding to CBC is prevented by ARS2. The different classifier factors compete for the same binding sites on both ARS2 and CBC in dynamic flux. Additional transcript-dependent cues then probably enable the binary interaction of PHAX or ZC3H18 with CBC. Given the partially overlapping binding sites of ARS2 and PHAX on CBC, in the case no RNA anomaly is detected, this might eventually result in exclusion of ARS2 favouring the assembly of the snRNA export complex. Alternatively, the binding of ZC3H18 to both ARS2 and CBC will result in recruitment of the NEXT complex and transcript degradation.

Concerning binary interactions with ARS2, the ARM, or ‘EDGEI’ motif, which binds to the effector domain of ARS2, has now been shown to be functionally important for a number of effectors (Dobrev et al., 2021; Foucher et al., 2022; Kiriyama et al., 2009; Melko et al., 2020; Polak et al., 2023; Rouviere et al., 2023; Schulze et al., 2018). Here, we identify a non-canonical, but functional, ARM at the N-terminus of human PHAX. We also report quantitative *in vitro* measurements of the affinity to ARS2 of seven different single ARMs occurring in productive factors and four different double or triple tandem ARMs found in degradative factors, showing that the latter are generally tighter binders. In the case of ZC3H18, which has a triple tandem ARM, based on an Alphafold prediction, we also show that the downstream Zn-knuckle provides additional interaction to ARS2.

Since ARS2 binds directly to the CBC and the considered factors (PHAX, NCBP3, ZC3H18) bind to both CBC and ARS2, one might expect that each factor can make a stable ternary complex with all binary interactions satisfied, i.e. a ‘triangular’ complex. We find that, *in vitro*, such ternary complexes are indeed formed but surprisingly in the presence of ARS2, these binary CBC-factor interactions are weakened, or lost, and the ternary complex is in fact a ‘linear’ CBC-ARS2-factor complex, with ARS2 responsible for recruitment of the factor. Thus, for instance disruption of the ARM of PHAX by a point mutation (E9R) is sufficient to release PHAX from the ternary complex, and similarly for ZC3H18 (Figure 5). This implies that ARS2 must bind to the CBC in a way that not only occludes the CBP20-CBP80 groove but also the Trp-helix binding site. Our structures of the CBC-ARS2 complexes provide an explanation for this, by showing that additional residues of ARS2, apart from those that bind in the CBP20-CBP80 groove indeed bind to the CBC, likely inhibiting Trp-helix binding. On the other hand, it should be kept in mind that this is a dynamic system with all interactions being of comparable affinity. Therefore, the linear CBC-factor-ARS2 complex is also possible, suggesting a mechanism by which multiple binary interactions can enhance the stability of a ternary complex, even if the triangular complex is not possible.

To clarify the importance of these interactions *in vivo*, we made disruptive mutations in the Trp-helix (W301E) and ARM of ZC3H18 and studied their behaviour in cells where endogenous ZC3H18 could be rapidly depleted (Figure 6). In interaction studies, we found, as *in vitro*, that abrogation of the ZC3H18-ARS2 interaction prevented co-immunoprecipitation of the CBC with ZC3H18. However, at the same time, the W301E mutation reduced the stability of the ternary complex. Importantly, neither mutation was capable of supporting ZC3H18-mediated degradation of several known targets of ZC3H18-NEXT. Thus, both binary interactions of ZC3H18 to the CBC and to ARS2 are functionally important, suggesting that at the point of irreversible fate decision, exclusion of other factors and other outcomes through competition at both sites is required. Based on this result, we propose that in the case of PHAX, at some point in the snRNA transcription process, additional transcript-dependent cues reinforce the binary interaction of PHAX with the CBC, perhaps with the competitive exclusion of ARS2 (consistent with the partially overlapping binding sites of ARS2 and PHAX on CBC), thus favouring the assembly of the snRNA export complex (Figure 7C). Further studies are required to reveal the exact cues that determine such transcript fate.

## Acknowledgements

SC and JK were funded by ANR MTREC (ANR-21-CE11-0021). THJ was supported by the Danish National Research Council and the Novo Nordisk Foundation [ExoAdapt Grant 31199]. IBS acknowledges integration into the Interdisciplinary Research Institute of Grenoble (IRIG, CEA). This work used the platforms of the Grenoble Instruct-ERIC center (ISBG ; UAR 3518 CNRS-CEA-UGA-EMBL) within the Grenoble Partnership for Structural Biology (PSB), supported by FRISBI (ANR-10-INBS-0005-02) and GRAL, financed within the University Grenoble Alpes graduate school (Ecoles Universitaires de Recherche) CBH-EUR-GS (ANR-17-EURE-0003). We thank Caroline Mas for assistance with ITC and Jean-Baptiste Reiser for his help and assistance with SPR and BLI. We thank Sarah Schneider and Romain Linares for support in using the EM Facility at EMBL Grenoble. We thank the European Synchrotron Radiation Facility for providing beam time on CM01 and Michael Hons and Romain Linares for their assistance. We thank Nadia Laurs Schmidt and Dorthe Caroline Riishøj for excellent technical assistance.

## Author contributions

E.D. with help of A.-E. F., K.H. and J.K performed all *in vitro* biochemical and biophysical characterization and cryo-EM sample preparation. E.P. and S.C. performed cryo-EM structural characterization of the CBC-NCBP3 complex, which was a pioneering achievement for the study. E.D. and S.C. performed cryo-EM structural characterization of CBC-PHAX and CBC-ARS2 complexes. MFB, WG and THJ performed all the cellular assays. S.C and J.K conceived the project and wrote the manuscript with input from all authors.

## Methods

### Protein Expression and Purification

Human CBC was reconstituted as described previously (Schulze and Cusack, 2017). Briefly, CBP80ΔNLS (lacking the N-terminal 19 residues) was expressed in High Five insect cells. CBP20 was expressed in *E. coli* BL21Star (DE3, Invitrogen) from the pETM30 vector (EMBL) as His-GST fusion. The cell pellets were lysed together in a buffer containing 20mM Tris pH 8, 100 mM NaCl and 2% glycerol. Lysates were clarified by centrifugation and the complex was purified on a Ni^2+^-Chelating Sepharose (Cytiva). The His-GST tag was cleaved off by TEV protease. Following a subsequent passage through a Ni^2+^-Chelating Sepharose, CBC was loaded on a Heparin HiTrap column (Cytiva) and further purified by a gel filtration on Superdex 200 (Cytiva).

Human ARS2^147-871^, FL-PHAX and its W118E and E9R mutants were expressed as His-tag fusions in *E. coli* BL21Star (DE3, Invitrogen) from the pETM11 vector (EMBL). The proteins were purified on a Ni^2+^-chelating Sepharose (Cytiva) followed by Hitrap Heparin column (Cytiva) and gel filtration on Superdex 200 (Cytiva) in a buffer containing 20mM Tris pH 8, 100 mM NaCl, 2% glycerol.

Human NCBP3, PHAX, ZC3H18 domains and their variants were cloned as His-MBP fusions into pETM41 vector (EMBL). Human NELF-E^244-380^ and its W345E mutant were expressed as His-tag fusions from pETM11 vector (EMBL). The proteins were expressed in *E. coli* BL21Star (DE3, Invitrogen) and purified on a Ni^2+^-chelating Sepharose (Cytiva) followed by gel filtration on Superdex 200 (Cytiva) in a buffer containing 20mM Tris pH 8, 100 mM NaCl, 2% glycerol.

### Assembly of CBC complexes

In order to prepare the sample for cryo-EM analysis, CBP80ΔNLS and His-GST-CBP20, His-MBP-NCBP3^560-620^, FL-PHAX or ARS2^147-871^ were purified separately. The His-MBP or His-GST fusion tags were cleaved off by TEV protease. Proteins were mixed together, the mixtures being supplemented by excess of cap analogue, cap-RNA or m^7^GTP. CBC-NCBP3^560-620^ complex was purified by an additional passage through a Ni^2+^-chelating Sepharose. Complexes were further purified by gel filtration on Superdex 200 (Cytiva).

### Pull-down assays

Purified CBC or CBC-ARS2^147-871^ and their mixtures with different His-MBP-tagged partners were loaded onto Amylose resin columns (NEB). Columns were then extensively washed with a buffer containing 100 mM Tris pH 8, 100 mM NaCl and 2 % glycerol. Bound proteins were eluted by addition of 10 mM maltose and analysed on SDS-PAGE.

### Isothermal Calorimetry (ITC)

ITC experiments were performed at 25°C using a PEAQ-ITC micro-calorimeter (MicroCal). Experiments included one 0.5 µl injection and 15-25 injections of 1.5-2.5 µL at 0.2-0.25mM of FL-PHAX or His-MBP-NCBP3, His-MBP-PHAX, His-MBP-ZC3H18 domains and their respective variants, into the sample cell that contained 20-30 µM CBC or ARS2^147-871^ in 20 mM Tris pH 8.0, 100 mM NaCl and 2 % glycerol. The initial data point was deleted from the data sets. Binding isotherms were fitted with a one-site binding model by nonlinear regression using the MicroCal PEAQ-ITC Analysis software.

### Biolayer interferometry (BLI)

Biolayer interferometry binding data were collected using integrated Octet RED96e (ForteBio) and processed using the integrated software. Similar quantities of biotinylated peptides were immobilized on streptavidin-coated biosensors (ForteBio, Menlo Park, CA). Peptides were diluted at concentrations of 0.1-1 µg/ml in TBS complemented with 0.02% Tween-20 as analysis buffer. Biosensors were pre-wetted in analysis buffer for 10 min, allowed to bind for 60-300s and washed in 10mM glycine. After equilibration in analysis buffer for 120 s, association and dissociation phases were monitored by dipping the functionalized biosensors in ARS2^147-871^ analyte solutions followed by analysis buffer for 160s each. Biosensor were regenerated using 10 mM glycine and equilibrated before each cycle. Signal from zero-concentration sample was subtracted from signals recorded for each analyte concentration. Specific signals at equilibrium were fitted using a Steady-state affinity determination method and a 1:1 interaction model.

### Surface plasmon resonance (SPR)

Direct binding studies were carried out in real-time on a Biacore T200 instrument (Biacore, Uppsala, Sweden). Similar levels of His-MBP-ZC3H18^298-339^ constructs were immobilized on independent channels on an NTA Sensor Chip (Cytiva) with a Ni^2+^ coated channel as a negative control. Binding assays were performed at 25 °C with a running buffer containing 100 mM NaCl, 20 mM Tris pH 8, 2% glycerol, complemented with 0.05% Tween-20 and 50µM EDTA. Indicated CBC complex concentration were injected at 30 µl per min for 100s. Proteins were allowed to dissociate for 200s at 30 µl per min. After each step, surface was regenerated using 10 mM NaOH and 350 mM EDTA. The kinetic rate constants K_D_, K_a_ and K_d_ were derived using Biacore T200 Evaluation Software by fitting the association and dissociation phases to a simple bimolecular interaction model.

### Cryo-EM specimen preparation

Before vitrification, cap analogue m^7^GpppG, m^7^Gppp-AAUCUAUAAUAG or m^7^GTP were added to each complex (at 20 µM) and the samples were left on ice for at least 1h. Cryo-EM specimens were prepared on Au300 R1.2/1.3 (Quantifoil) that were glow-discharged for 45s at 25 mA (PELCO easy glow). A drop of 3.5-4 µl sample was applied on one side in a Vitrobot (Vitrobot Mk IV, Thermo Fisher Scientific) at 4 °C and 100 % humidity. Blotting was run at force 0, for a total time of 2-3 s. Grids were then vitrified by plunging into liquid ethane at liquid N2 temperature. Grids were clipped into autoloader cartridges and screened using a Glacios cryo-electron microscope (Thermo Fisher Scientific) equipped with a Falcon 4 detector (EMBL Grenoble). Promising samples, corresponding to well-distributed particles being visible by eye at -1 µM defocus, were used for data collection at CM01, ESRF Grenoble France.

### Cryo-EM Data Collection and Processing

The CBC-m^7^GpppG-NCBP3^560-620^dataset was acquired using Titan Krios CM01 (ESRF, Grenoble, France)(Kandiah et al., 2019) operating at 300 keV, equipped with K2 Quantum detector (Gatan) and a GIF Quantum energy filter (Gatan). The CBC-capRNA- PHAX and CBC-m^7^GTP-ARS2^147-871^ datasets were acquired on the same microscope but equipped with a K3 Quantum detector camera (Gatan).

For the CBC-m^7^GpppG-NCBP3^560-620^ complex, 9629 movies were collected at 165K magnification, corresponding to a pixel size of 0.827 px/Å for a total dose of 40 e-/A^2^. For the CBC-capRNA-PHAX complex, 6517 movies were collected at 165K magnification, with a pixel size of 0.84 px/Å and a total dose of 41.7 e-/A^2^. For the CBC-m^7^GTP-ARS2^147-871^, 12228 movies were collected at 165K magnification, with a pixel size of 0.84 px/Å and a total dose of 40 e-/A^2^. For all datasets, movies were imported into CryoSPARC (Punjani et al., 2017) and aligned and dose-weighted using Patch MotionCorr. CTF estimation was performed with Patch CTF and micrographs were manually curated according to CTF based estimated resolution (<10Å) and estimated ice thickness (<1.1).

Particles were firstly picked on 100-200 representative micrographs using Blob Picker or Topaz (Bepler et al., 2019) and then extracted with a box size of 300×300 pixels (NCBP3) or 256×256 (PHAX and ARS2 datasets) (Figure S2). The best particles were then used to generate a Topaz model or to create Templates for particles picking in the remaining micrographs. After extraction, 2D classification was applied to eliminate bad particles and to select 2D class averages with lower background noise and stronger features. One to two cycles of *ab-initio* reconstruction were then performed, followed by heterogeneous refinement. For the PHAX and ARS2 containing complexes, 3D variability analysis was performed to further select particles (Punjani and Fleet, 2021). Importantly, the coordination of CBC to its partner always correlates with full structuring of CBP20, including its N-terminal region, a hallmark of cap-binding (Mazza et al., 2002)(Figure S2G). Non-uniform (NU) refinement was finally performed on the final sets of particles.

The final map was at an average resolution of 3.19 Å for CBC-m^7^GpppG-NCBP3^560-620^, 3.30 Å for CBC-capRNA-PHAX and 3.40 Å for the CBC-m^7^GTP-ARS2^147-871^ (FSC 0.143 threshold) (Table S1).

All final maps were used to calculate directional FSC and local resolution in CryoSPARC. For model building in COOT, the map was sharpened/blurred using the mrc_to_mtz tool in CCPEM (Burnley et al., 2017). To make figures in ChimeraX 1.4 (Pettersen et al., 2021), maps were locally filtered based on resolution and sharpened (see Table S1). Processing workflows, comprising data statistics, are shown in Figures S2, S6 and S11.

### Model building and refinement

Models were iteratively constructed using COOT (Emsley and Cowtan, 2004) and PHENIX real-space refinement (Afonine et al., 2018) starting with the published CBC- mGTP-ASR2 structure (PDB: 5OO6). Refinement and validation statistics, obtained using the PHENIX validation tool are reported in Table S1. For the CBC-capRNA-PHAX and CBC-m^7^GTP-ARS2^147-871^, model building was guided partly by the AlphaFold model (Figure S6), which correctly predicted the PHAX Trp118 helix and extreme C-terminus binding. There is no clear density for the capped RNA apart from the m^7^GTP although there is unassignable density near the cap-binding site that could be RNA or another part of PHAX. Additional extra, but disjointed density close to CBP20 Y142 and Y149 and between CBP20 Y50 and CBP80 Y461 (the conserved arginine binding site for ARS2 and NELF-E) could not be interpreted.

Software used in this project was installed and configured by SBGrid (Morin et al., 2013).

### Cell culture and transfections

All cell lines used or generated in this study were descendants of E14TG2a mouse embryonic stem (mES) cells (male genotype, XY). mES cells were cultured on 0.2% gelatin coated plates in 2i/LIF containing medium (1:1 mix of DMEM/F12 (Gibco) and Neurobasal (Gibco) supplemented with 1x Pen-Strep (Gibco), 2 µM Glutamax, 50 µM beta-mercaptoethanol (Gibco), 0.1 mM Non-Essential Amino Acids (Gibco), 1 mM sodium pyruvate (Gibco), 0.5x N2 Supplement (Gibco), 0.5x B27 Supplement (Gibco), 3 µM GSK3- inhibitor (CHIR99021), 1 µM MEK-inhibitor (PD0325901) and Leukemia Inhibitory Factor (LIF, produced in house) at 37°C, 5% CO_2_. Cells were passaged every 48-72 hours by aspirating medium, dissociating cells with 0.05% Trypsin-EDTA (Gibco) briefly at 37°C before neutralizing with an equal volume of 1x Trypsin Inhibitor (Sigma) and gentle disruption by pipetting. Cells were pelleted by centrifugation to remove Trypsin before resuspending in 2i/LIF medium and plating ∼ 8×10^4^ cells/ml. Cell lines were transfected with plasmids using Viafect (Promega) in 6 well plates according to the manufacturer’s instructions. For antibiotic selection, Blasticidin (BSD) was used at 10 µg/ml. For depletions in mAID-tagged cell lines, 750 µM indole-3-acetic acid sodium salt (IAA, Sigma) was supplemented to the medium and cells were incubated for the indicated time points before harvest.

### CRISPR/Cas9 knock-in cell lines

CRISPR/Cas9 mediated genomic knock-ins of C-terminal 3xFLAG(3F) mini-AID(mAID) tags were carried out using pGolden (pGCT) homology dependent repair (HDR) donor vectors (Garland et al., 2022; Gockert et al., 2022). The generation of *Zc3h18-3F- mAID* cell lines was described in (Polak et al., 2023). Single guide RNAs (sgRNAs) were designed using the CHOPCHOP tool (v3) (Labun et al., 2019) and cloned into the pSLCas(BB)2A-PURO vector (pX459 Addgene plasmid ID: #48139) as previously described (Ran et al., 2013). OsTIR1-HA expressing mES cells (Garland et al., 2022) were co-transfected using 2 HDR donor vectors harboring distinct selection markers (HYG/NEO) along with a sgRNA/Cas9 vector in a 1:1:1 ratio. Colonies were maintained under HYG/NEO double selection for the donor plasmid markers to increase the likelihood of homozygous knock-in cells. Single cell clones, that survived the selection process, were expanded before screening by western blotting analysis and confirmed by Sanger sequencing of the target locus. All cell lines used and generated in this study are listed in Table S2.

### cDNA cloning and exogenous expression of ZC3H18

Full-length mouse ZC3H18 cDNA (NM_001029993) was cloned from cDNA libraries synthesized from 2 µg total RNA using TaqMan Reverse Transcription reagents (Thermo) into a pCR8/GW/TOPO Gateway entry plasmid (Thermo) by TA cloning. The pCR8[mZC3H18] plasmid was used as a template to generate subsequent ZC3H18 cDNA constructs that were cloned into a piggyBAC (pB) vector containing an N-terminal MYC tag and BSD selection marker using NEBbuilder HiFI DNA assembly (NEB). *OsTIR1-HA, Zc3h18-3F-mAID* mES cells were transfected with pB-MYC-mZC3H18^x^-BSD vectors along with a piggyBAC transposase expressing vector (pBase) in a 1:1 ratio using Viafect (Promega). Cell pools were selected with BSD for ∼ 7-10 days or until negative control cells no longer survived. Expression of constructs were validated by western blotting analysis using MYC antibodies. All generated constructs used and generated in this study are listed in Table S3.

### RNA isolation and RT-qPCR analysis

Total RNA was isolated from cells using TRIzol reagent (Invitrogen) according to the manufacturer’s instructions. Extracted RNA was treated with TURBO DNase (Invitrogen) according to the manufacturer’s instructions, followed by cDNA synthesis from 2 µg RNA using SuperScript III Reverse Transcriptase (Invitrogen) and a mixture of 80 pmol random primers (Invitrogen) and 20 pmol oligo d(T)_20_VN (Merck) primers. qPCR was performed using Platinum SYBR Green (Invitrogen) and AriaMX Real-Time PCR machine (Agilent Technologies). Primers used for qPCR are listed in Table S4.

### Western blotting analysis

Whole cell protein lysates were prepared using RSB100 lysis buffer (10 mM Tris-HCl pH 7.5, 100 mM NaCl, 2.5 mM MgCl_2_, 0.5% NP-40, 0.5% Triton X-100) freshly supplemented with protease inhibitors (Roche). Samples were denatured by the addition of NuPAGE loading buffer (Invitrogen) and NuPAGE Sample Reducing Agent (Invitrogen) before boiling at 95°C for 5 minutes. SDS-PAGE was carried out on either NuPAGE 4-12% Bis-Tris or 3-8% Tris-Acetate gels (Invitrogen). Western blotting analysis was carried out according to standard protocols with the antibodies listed in Table S5.

### Immunoprecipitation (IP) experiments

Whole cell lysates from ∼ 2×10^7^ cells were extracted in HT150 extraction buffer (20 mM HEPES pH 7.4, 150 mM NaCl, 0.5% Triton X-100) freshly supplemented with protease inhibitors. Lysates were sheared by sonication (3 × 5s, amplitude 2) using a Sonifier 450 (Branson) and cleared by centrifugation at 18,000 rcf for 20 minutes. Clarified lysates were incubated with Pierce anti-c-MYC magnetic beads (Thermo) overnight at 4°C. Beads were washed 3x with HT150 extraction buffer and transferred to a fresh tube on the final wash. Proteins were eluted by incubation at 95°C in NuPAGE Loading Buffer (Invitrogen) for 5 minutes. Eluates were mixed with NuPAGE Sample reducing agent (Invitrogen) and denatured for a further 5 minutes at 95°C before proceeding with western blotting analysis.

## References

Afonine, P.V., Poon, B.K., Read, R.J., Sobolev, O.V., Terwilliger, T.C., Urzhumtsev, A., and Adams, P.D. (2018). Real-space refinement in PHENIX for cryo-EM and crystallography. Acta Crystallogr D Struct Biol 74, 531–544.

Andersen, P.R., Domanski, M., Kristiansen, M.S., Storvall, H., Ntini, E., Verheggen, C., Schein, A., Bunkenborg, J., Poser, I., Hallais, M., et al. (2013). The human cap-binding complex is functionally connected to the nuclear RNA exosome. Nature Structural & Molecular Biology 20, 1367–1376.

Aoi, Y., Smith, E.R., Shah, A.P., Rendleman, E.J., Marshall, S.A., Woodfin, A.R., Chen, F.X., Shiekhattar, R., and Shilatifard, A. (2020). NELF Regulates a Promoter-Proximal Step Distinct from RNA Pol II Pause-Release. Mol Cell 78, 261–274 e265.

Bepler, T., Morin, A., Rapp, M., Brasch, J., Shapiro, L., Noble, A.J., and Berger, B. (2019). Positive-unlabeled convolutional neural networks for particle picking in cryo-electron micrographs. Nat Methods 16, 1153–1160.

Boulon, S., Verheggen, C., Jady, B.E., Girard, C., Pescia, C., Paul, C., Ospina, J.K., Kiss, T., Matera, A.G., Bordonne, R., et al. (2004). PHAX and CRM1 are required sequentially to transport U3 snoRNA to nucleoli. Mol Cell 16, 777–787.

Burnley, T., Palmer, C.M., and Winn, M. (2017). Recent developments in the CCP-EM software suite. Acta Crystallogr D Struct Biol 73, 469–477.

Cheng, H., Dufu, K., Lee, C.S., Hsu, J.L., Dias, A., and Reed, R. (2006). Human mRNA export machinery recruited to the 5’ end of mRNA. Cell 127, 1389–1400.

Cho, H., Rambout, X., Gleghorn, M.L., Nguyen, P.Q.T., Phipps, C.R., Miyoshi, K., Myers, J.R., Kataoka, N., Fasan, R., and Maquat, L.E. (2018). Transcriptional coactivator PGC- 1alpha contains a novel CBP80-binding motif that orchestrates efficient target gene expression. Genes Dev 32, 555–567.

Cordiner, R.A., Dou, Y., Thomsen, R., Bugai, A., Granneman, S., and Heick Jensen, T. (2023). Temporal-iCLIP captures co-transcriptional RNA-protein interactions. Nat Commun 14, 696.

Dantsuji, S., Ohno, M., and Taniguchi, I. (2023). The hnRNP C tetramer binds to CBC on mRNA and impedes PHAX recruitment for the classification of RNA polymerase II transcripts. Nucleic Acids Res.

Ding, S., Wu, X., Li, G., Han, M., Zhuang, Y., and Xu, T. (2005). Efficient transposition of the piggyBac (PB) transposon in mammalian cells and mice. Cell 122, 473–483.

Dobrev, N., Ahmed, Y.L., Sivadas, A., Soni, K., Fischer, T., and Sinning, I. (2021). The zinc-finger protein Red1 orchestrates MTREC submodules and binds the Mtl1 helicase arch domain. Nat Commun 12, 3456.

Dou, Y., Barbosa, I., Jiang, H., Iasillo, C., Molloy, K.R., Schulze, W.M., Cusack, S., Schmid, M., Le Hir, H., LaCava, J., et al. (2020). NCBP3 positively impacts mRNA biogenesis. Nucleic Acids Res 48, 10413–10427.

Emsley, P., and Cowtan, K. (2004). Coot: model-building tools for molecular graphics. Acta Crystallogr D Biol Crystallogr 60, 2126–2132.

Evans, R., O’Neill, M., Pritzel, A., Antropova, N., Senior, A., Green, T., Žídek, A., Bates, R., Blackwell, S., Yim, J., et al. (2022). Protein complex prediction with AlphaFold-Multimer. bioRxiv, 2021.2010.2004.463034.

Fan, J., Kuai, B., Wu, G., Wu, X., Chi, B., Wang, L., Wang, K., Shi, Z., Zhang, H., Chen, S., et al. (2017). Exosome cofactor hMTR4 competes with export adaptor ALYREF to ensure balanced nuclear RNA pools for degradation and export. EMBO J 36, 2870–2886.

Foucher, A.E., Touat-Todeschini, L., Juarez-Martinez, A.B., Rakitch, A., Laroussi, H., Karczewski, C., Acajjaoui, S., Soler-Lopez, M., Cusack, S., Mackereth, C.D., et al. (2022). Structural analysis of Red1 as a conserved scaffold of the RNA-targeting MTREC/PAXT complex. Nat Commun 13, 4969.

Gable, D.L., Gaysinskaya, V., Atik, C.C., Talbot, C.C., Jr., Kang, B., Stanley, S.E., Pugh, E.W., Amat-Codina, N., Schenk, K.M., Arcasoy, M.O., et al. (2019). ZCCHC8, the nuclear exosome targeting component, is mutated in familial pulmonary fibrosis and is required for telomerase RNA maturation. Genes Dev 33, 1381–1396.

Garland, W., and Jensen, T.H. (2020). Nuclear sorting of RNA. Wiley Interdiscip Rev RNA 11, e1572.

Garland, W., Muller, I., Wu, M., Schmid, M., Imamura, K., Rib, L., Sandelin, A., Helin, K., and Jensen, T.H. (2022). Chromatin modifier HUSH co-operates with RNA decay factor NEXT to restrict transposable element expression. Mol Cell 82, 1691–1707 e1698.

Gebhardt, A., Bergant, V., Schnepf, D., Moser, M., Meiler, A., Togbe, D., Mackowiak, C., Reinert, L.S., Paludan, S.R., Ryffel, B., et al. (2019). The alternative cap-binding complex is required for antiviral defense in vivo. PLoS Pathog 15, e1008155.

Gebhardt, A., Habjan, M., Benda, C., Meiler, A., Haas, D.A., Hein, M.Y., Mann, A., Mann, M., Habermann, B., and Pichlmair, A. (2015). mRNA export through an additional cap-binding complex consisting of NCBP1 and NCBP3. Nat Commun 6, 8192.

Gerlach, P., Garland, W., Lingaraju, M., Salerno-Kochan, A., Bonneau, F., Basquin, J., Jensen, T.H., and Conti, E. (2022). Structure and regulation of the nuclear exosome targeting complex guides RNA substrates to the exosome. Mol Cell 82, 2505–2518 e2507.

Giacometti, S., Benbahouche, N.E.H., Domanski, M., Robert, M.C., Meola, N., Lubas, M., Bukenborg, J., Andersen, J.S., Schulze, W.M., Verheggen, C., et al. (2017). Mutually Exclusive CBC-Containing Complexes Contribute to RNA Fate. Cell Rep 18, 2635–2650.

Gockert, M., Schmid, M., Jakobsen, L., Jens, M., Andersen, J.S., and Jensen, T.H. (2022). Rapid factor depletion highlights intricacies of nucleoplasmic RNA degradation. Nucleic Acids Res 50, 1583–1600.

Gromadzka, A.M., Steckelberg, A.L., Singh, K.K., Hofmann, K., and Gehring, N.H. (2016). A short conserved motif in ALYREF directs cap- and EJC-dependent assembly of export complexes on spliced mRNAs. Nucleic Acids Res 44, 2348–2361.

Hallais, M., Pontvianne, F., Andersen, P.R., Clerici, M., Lener, D., Benbahouche, N.E., Gostan, T., Vandermoere, F., Robert, M.C., Cusack, S., et al. (2013). CBC-ARS2 stimulates 3 ’-end maturation of multiple RNA families and favors cap-proximal processing. Nature Structural & Molecular Biology 20, 1358–1366.

Iasillo, C., Schmid, M., Yahia, Y., Maqbool, M.A., Descostes, N., Karadoulama, E., Bertrand, E., Andrau, J.C., and Jensen, T.H. (2017). ARS2 is a general suppressor of pervasive transcription. Nucleic Acids Res 45, 10229–10241.

Jumper, J., Evans, R., Pritzel, A., Green, T., Figurnov, M., Ronneberger, O., Tunyasuvunakool, K., Bates, R., Zidek, A., Potapenko, A., et al. (2021). Highly accurate protein structure prediction with AlphaFold. Nature 596, 583–589.

Kandiah, E., Giraud, T., de Maria Antolinos, A., Dobias, F., Effantin, G., Flot, D., Hons, M., Schoehn, G., Susini, J., Svensson, O., et al. (2019). CM01: a facility for cryo-electron microscopy at the European Synchrotron. Acta Crystallogr D Struct Biol 75, 528–535.

Kiriyama, M., Kobayashi, Y., Saito, M., Ishikawa, F., and Yonehara, S. (2009). Interaction of FLASH with arsenite resistance protein 2 is involved in cell cycle progression at S phase. Mol Cell Biol 29, 4729–4741.

Labun, K., Montague, T.G., Krause, M., Torres Cleuren, Y.N., Tjeldnes, H., and Valen, E. (2019). CHOPCHOP v3: expanding the CRISPR web toolbox beyond genome editing. Nucleic Acids Res 47, W171–W174.

Lubas, M., Christensen, M.S., Kristiansen, M.S., Domanski, M., Falkenby, L.G., Lykke-Andersen, S., Andersen, J.S., Dziembowski, A., and Jensen, T.H. (2011). Interaction profiling identifies the human nuclear exosome targeting complex. Mol Cell 43, 624–637.

Lykke-Andersen, S., Rouviere, J.O., and Jensen, T.H. (2021). ARS2/SRRT: at the nexus of RNA polymerase II transcription, transcript maturation and quality control. Biochem Soc Trans 49, 1325–1336.

Mazza, C., Segref, A., Mattaj, I.W., and Cusack, S. (2002). Large-scale induced fit recognition of an m(7)GpppG cap analogue by the human nuclear cap-binding complex. Embo Journal 21, 5548–5557.

McCloskey, A., Taniguchi, I., Shinmyozu, K., and Ohno, M. (2012). hnRNP C tetramer measures RNA length to classify RNA polymerase II transcripts for export. Science 335, 1643–1646.

Melko, M., Winczura, K., Rouviere, J.O., Oborska-Oplova, M., Andersen, P.K., and Heick Jensen, T. (2020). Mapping domains of ARS2 critical for its RNA decay capacity. Nucleic Acids Res 48, 6943–6953.

Meola, N., Domanski, M., Karadoulama, E., Chen, Y., Gentil, C., Pultz, D., Vitting-Seerup, K., Lykke-Andersen, S., Andersen, J.S., Sandelin, A., et al. (2016). Identification of a Nuclear Exosome Decay Pathway for Processed Transcripts. Mol Cell 64, 520–533.

Morin, A., Eisenbraun, B., Key, J., Sanschagrin, P.C., Timony, M.A., Ottaviano, M., and Sliz, P. (2013). Collaboration gets the most out of software. Elife 2, e01456.

Mourao, A., Varrot, A., Mackereth, C.D., Cusack, S., and Sattler, M. (2010). Structure and RNA recognition by the snRNA and snoRNA transport factor PHAX. Rna-a Publication of the Rna Society 16, 1205–1216.

Muller-McNicoll, M., and Neugebauer, K.M. (2014). Good cap/bad cap: how the cap-binding complex determines RNA fate. Nat Struct Mol Biol 21, 9–12.

Narita, T., Yung, T.M.C., Yamamoto, J., Tsuboi, Y., Tanabe, H., Tanaka, K., Yamaguchi, Y., and Handa, H. (2007). NELF interacts with CBC and participates in 3 ’ end processing of replication-dependent histone rnRNAs. Molecular Cell 26, 349–365.

Natsume, T., and Kanemaki, M.T. (2017). Conditional Degrons for Controlling Protein Expression at the Protein Level. Annu Rev Genet 51, 83–102.

O’Sullivan, C., Christie, J., Pienaar, M., Gambling, J., Nickerson, P.E.B., Alford, S.C., Chow, R.L., and Howard, P.L. (2015). Mutagenesis of ARS2 Domains To Assess Possible Roles in Cell Cycle Progression and MicroRNA and Replication-Dependent Histone mRNA Biogenesis. Molecular and Cellular Biology 35, 3753–3767.

Ohno, M., Segref, A., Bachi, A., Wilm, M., and Mattaj, I.W. (2000). PHAX, a mediator of U snRNA nuclear export whose activity is regulated by phosphorylation. Cell 101, 187–198.

Pabis, M., Neufeld, N., Steiner, M.C., Bojic, T., Shav-Tal, Y., and Neugebauer, K.M. (2013). The nuclear cap-binding complex interacts with the U4/U6.U5 tri-snRNP and promotes spliceosome assembly in mammalian cells. RNA 19, 1054–1063.

Pettersen, E.F., Goddard, T.D., Huang, C.C., Meng, E.C., Couch, G.S., Croll, T.I., Morris, J.H., and Ferrin, T.E. (2021). UCSF ChimeraX: Structure visualization for researchers, educators, and developers. Protein Sci 30, 70–82.

Polak, P., Garland, W., Schmid, M., Salerno-Kochan, A., Jakobsen, L., Bockert, M., Rathore, O., Gerlach, P., Silla, T., Andersen, J.-S., et al. (2023). Dual agonistic and antagonistic roles of ZC3H18 provides for co-activation of distinct nuclear RNA decay pathways. BioRxiv.

Punjani, A., and Fleet, D.J. (2021). 3D variability analysis: Resolving continuous flexibility and discrete heterogeneity from single particle cryo-EM. J Struct Biol 213, 107702.

Punjani, A., Rubinstein, J.L., Fleet, D.J., and Brubaker, M.A. (2017). cryoSPARC: algorithms for rapid unsupervised cryo-EM structure determination. Nat Methods 14, 290–296.

Puno, M.R., and Lima, C.D. (2022). Structural basis for RNA surveillance by the human nuclear exosome targeting (NEXT) complex. Cell 185, 2132–2147 e2126.

Rambout, X., Cho, H., Blanc, R., Lyu, Q., Miano, J.M., Chakkalakal, J.V., Nelson, G.M., Yalamanchili, H.K., Adelman, K., and Maquat, L.E. (2023). PGC-1alpha senses the CBC of pre-mRNA to dictate the fate of promoter-proximally paused RNAPII. Mol Cell 83, 186–202 e111.

Rambout, X., and Maquat, L.E. (2020). The nuclear cap-binding complex as choreographer of gene transcription and pre-mRNA processing. Genes Dev 34, 1113–1127.

Rambout, X., and Maquat, L.E. (2021). NCBP3: A Multifaceted Adaptive Regulator of Gene Expression. Trends Biochem Sci 46, 87–96.

Ran, F.A., Hsu, P.D., Wright, J., Agarwala, V., Scott, D.A., and Zhang, F. (2013). Genome engineering using the CRISPR-Cas9 system. Nat Protoc 8, 2281–2308.

Rouviere, J.O., Salerno-Kochan, A., Lykke-Andersen, S., Garland, W., Dou, Y., Rathore, O., Molska, E.S., Wu, G., Schmid, M., Bugai, A., et al. (2023). ARS2 instructs early termination-coupled RNA decay by recruiting ZC3H4 to nascent transcripts. Mol Cell. 83, 2240–2257e6.

Schulze, W.M., and Cusack, S. (2017). Structural basis for mutually exclusive co-transcriptional nuclear cap-binding complexes with either NELF-E or ARS2. Nat Commun 8, 1302.

Schulze, W.M., Stein, F., Rettel, M., Nanao, M., and Cusack, S. (2018). Structural analysis of human ARS2 as a platform for co-transcriptional RNA sorting. Nat Commun 9, 1701.

Segref, A., Mattaj, I.W., and Ohno, M. (2001). The evolutionarily conserved region of the U snRNA export mediator PHAX is a novel RNA-binding domain that is essential for U snRNA export. RNA 7, 351–360.

Shen, Z., Zeng, L., and Zhang, Z. (2020). Translatome and Transcriptome Profiling of Hypoxic-Induced Rat Cardiomyocytes. Mol Ther Nucleic Acids 22, 1016–1024.

Singh, G., Seufzer, B., Song, Z., Zucko, D., Heng, X., and Boris-Lawrie, K. (2022). HIV-1 hypermethylated guanosine cap licenses specialized translation unaffected by mTOR. Proc Natl Acad Sci U S A 119.

Suzuki, T., Izumi, H., and Ohno, M. (2010). Cajal body surveillance of U snRNA export complex assembly. J Cell Biol 190, 603–612.

Townsend, C., Leelaram, M.N., Agafonov, D.E., Dybkov, O., Will, C.L., Bertram, K., Urlaub, H., Kastner, B., Stark, H., and Luhrmann, R. (2020). Mechanism of protein-guided folding of the active site U2/U6 RNA during spliceosome activation. Science 370.

Viphakone, N., Sudbery, I., Griffith, L., Heath, C.G., Sims, D., and Wilson, S.A. (2019). Co-transcriptional Loading of RNA Export Factors Shapes the Human Transcriptome. Mol Cell 75, 310–323 e318.

Winczura, K., Schmid, M., Iasillo, C., Molloy, K.R., Harder, L.M., Andersen, J.S., LaCava, J., and Jensen, T.H. (2018). Characterizing ZC3H18, a Multi-domain Protein at the Interface of RNA Production and Destruction Decisions. Cell Rep 22, 44–58.

Zhou, Y., Zhu, J., Schermann, G., Ohle, C., Bendrin, K., Sugioka-Sugiyama, R., Sugiyama, T., and Fischer, T. (2015). The fission yeast MTREC complex targets CUTs and unspliced pre-mRNAs to the nuclear exosome. Nat Commun 6, 7050.

Zinder, J.C., and Lima, C.D. (2017). Targeting RNA for processing or destruction by the eukaryotic RNA exosome and its cofactors. Genes Dev 31, 88–100.

