## Supplementary information for "Structural basis for competitive binding of productive and degradative co-transcriptional effectors to the nuclear cap-binding complex"

Contains 11 Supplemental Figures, 5 Supplemental Tables and Supplemental references.

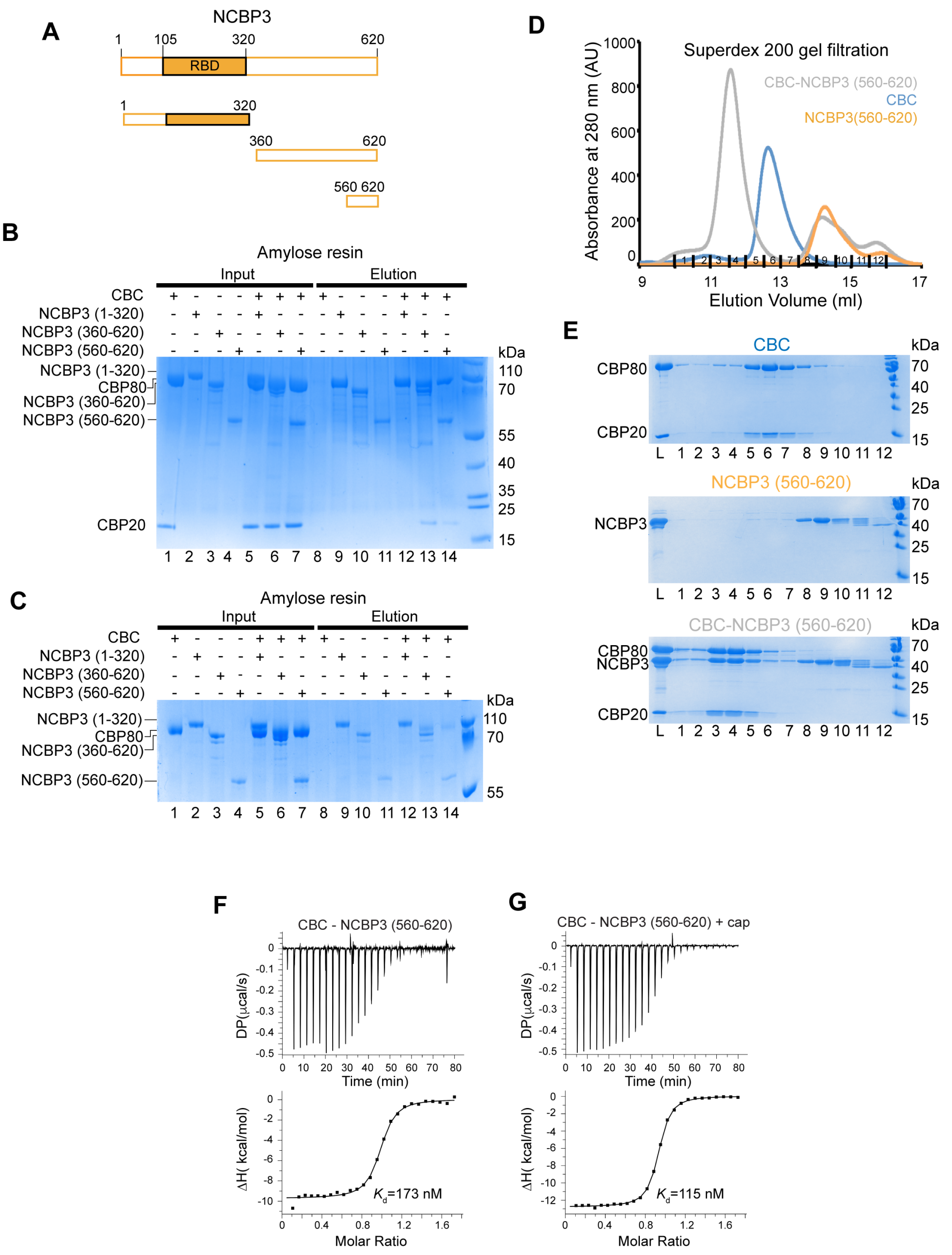


**Figure S1. Biochemical analysis of the CBC-NCBP3 interaction.**
**(A)** Schematic representation of the NCBP3 domain structure and constructs used in this study. RBD: RNA binding domain. **(B)** MBP pull-down experiments of His-MBP-tagged NCBP3 constructs with CBC in presence of cap analogue. All proteins were first purified by affinity chromatography and gel filtration. Proteins were mixed as indicated above the lanes. A total of 1% of the input (lanes 1–7) and 0.5% of the eluates (lanes 8–14) were analysed on 15% SDS-PAGE gels stained with Commassie brilliant blue. CBC is not retained by NCBP3$^{1\text{-}320}$ (lane 12), whereas the C-terminal constructs NCBP3$^{360\text{-}620}$ and NCBP3$^{560\text{-}620}$ are sufficient for the interaction (lanes 13 and 14). **(C)** To better distinguish the NCBP3 and CBP80 bands of similar size, a 7% SDS-PAGE gel of the MBP pull-down assays shown in **B** was performed**.** **(D)** Overlay of Superdex 200 gel filtration elution profiles of CBC, His-MBP-NCBP3$^{560\text{-}620}$ and their complex. **(E)** SDS-PAGE analysis of fractions 1-12 of the Superdex 200 gel filtration elution profiles shown in **D**. **(F, G)** ITC measurements of the interaction affinity between His-MBP-NCBP3$^{560\text{-}620}$ and CBC in the absence or presence of cap analogue.

**Cryo-EM structure determination of CBC-m$^7$GpppG-NCBP3$^{560-620}$**

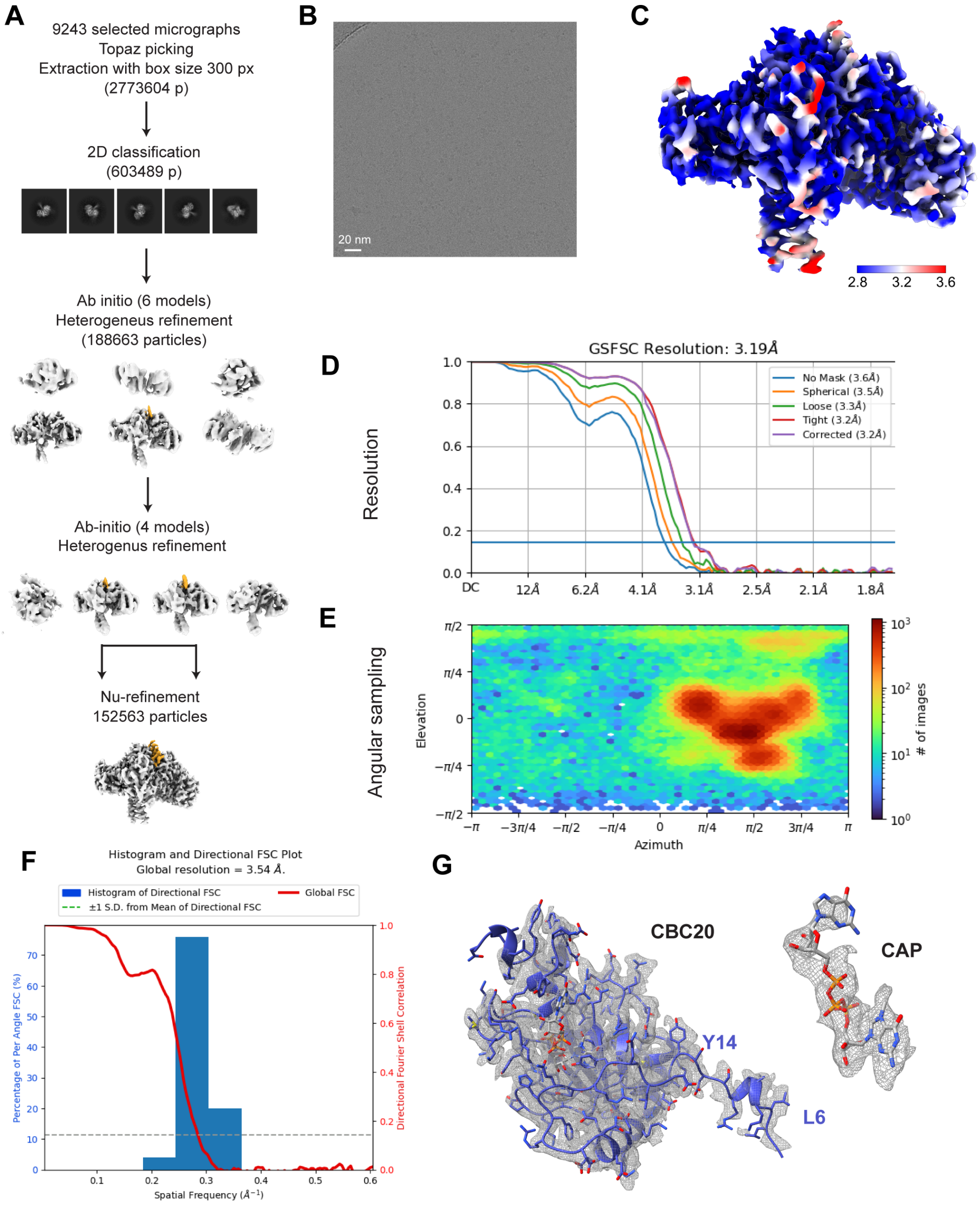


**Figure S2. Cryo-EM analysis of the CBC-m$^7$GpppG-NCBP3$^{560-620}$ complex.**
**(A)** Summary of the cryo-EM processing workflow used to obtain the final map, as described in the Methods section. The cryo-EM maps are unsharpened and the location of NCBP3 peptide is shown in gold. **(B)** Example of an aligned micrograph. **(C)** Final cryo-EM density filtered based on the resolution, sharpened with B-factor -100Å$^2$ and colored according to local resolution. **(D)** Map resolution according to the FSC threshold of 0.143. **(E)** Angular sampling distribution scheme. **(F)** Map resolution according to the 3D FSC. **(G)** Cryo-EM density corresponding to CBP20 and m$^7$GpppG.

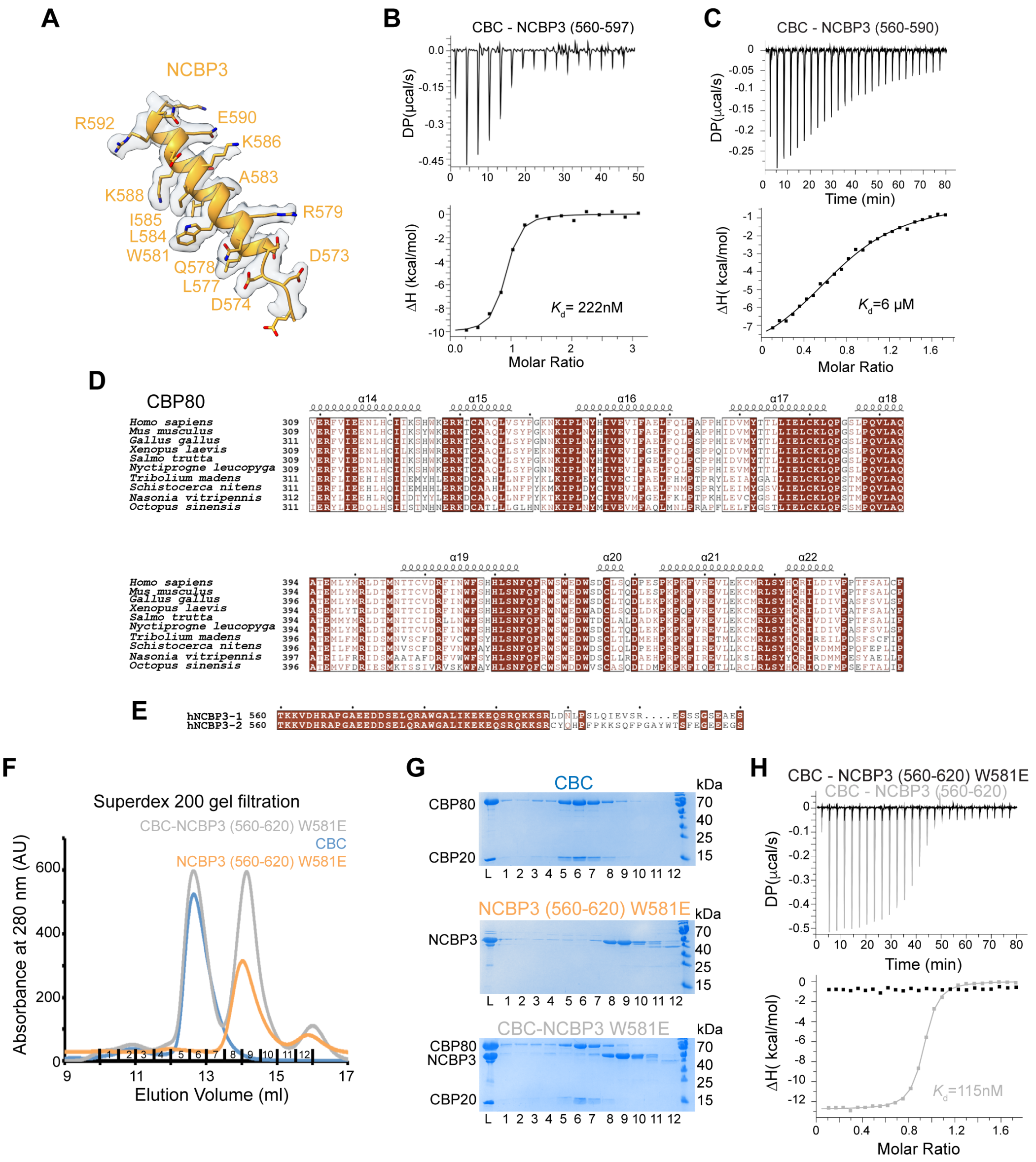


**Figure S3. Further details of the CBC-NCBP3 interaction.**
**(A)** Cryo-EM density for the NCBP3$^{560\text{-}620}$ peptide within the CBC-m$^7$GpppG-NCBP3$^{560\text{-}620}$ complex**.** **(B, C)** ITC measurements of the interaction affinity between CBC and His-MBP-NCBP3$^{560\text{-}597}$ or His-MBP-NCBP3$^{560\text{-}590}$, respectively, in the presence of cap analogue. **(D)** Sequence alignment of representative CBP80 proteins. Only the sequence of residues 309-478 covering the second MIF4G domain is shown. Identical residues are in brown boxes. **(E)** Sequence alignment of human NCBP3 isoform 1 and 2. Isoform 2 differs from isoform 1 after residue 597. **(F)** Overlay of Superdex 200 gel filtration elution profiles of CBC, NCBP3$^{560\text{-}620}$ (W581E) and their mixture. **(G)** SDS-PAGE analysis of the Superdex 200 gel filtration elution profiles shown in **F**. L indicates input samples loaded onto the column. Compare with Supplementary Fig. 1d,e. **(H)** ITC measurements of the interaction affinity between CBC and WT (grey) or W581E mutant (black) of His-MBP-NCBP3$^{560\text{-}620}$ in the presence of cap analogue.

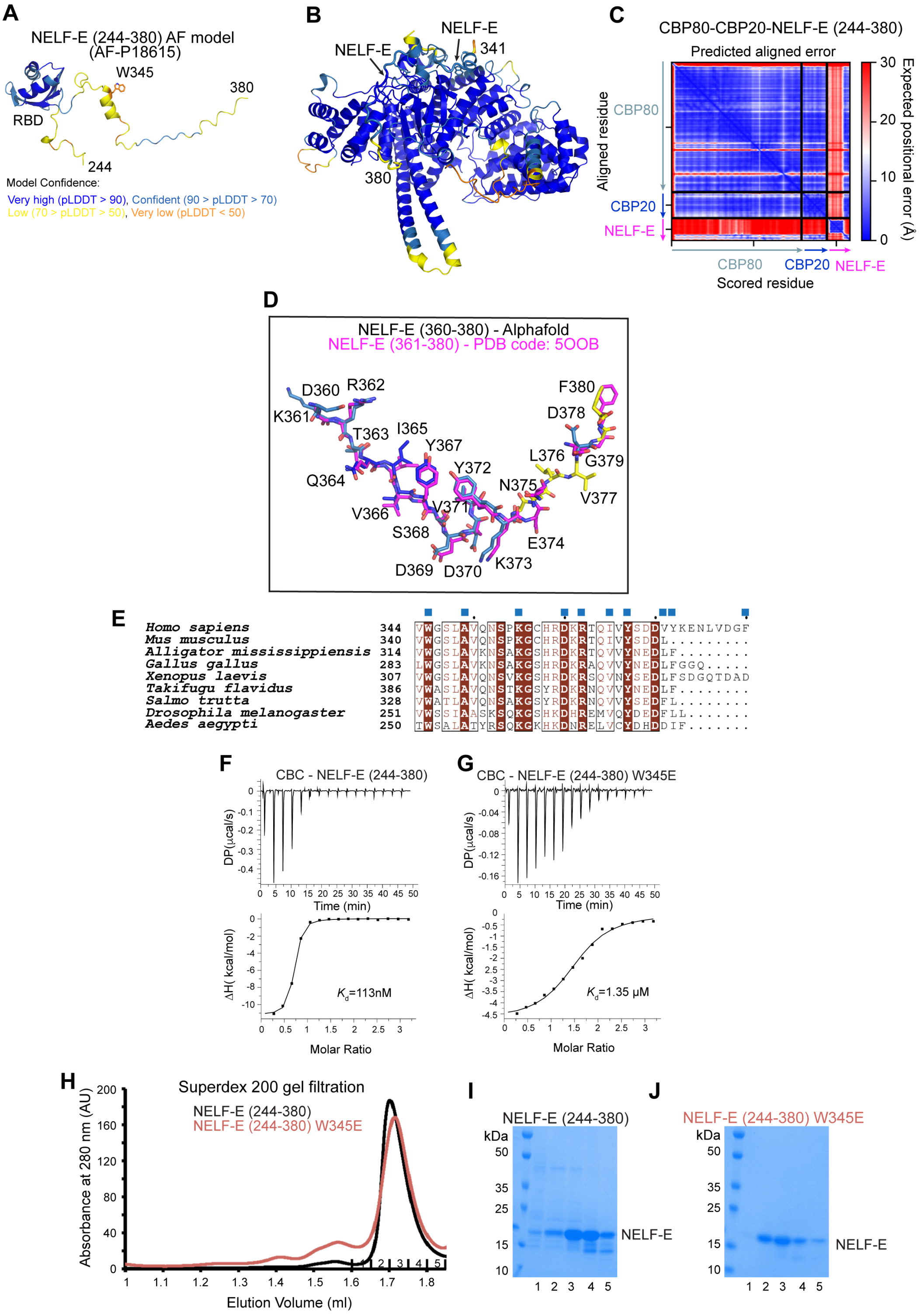


**Figure S4. NELF-E interaction with CBC.**
**(A)** The Alphafold database model of NELF-E$^{244\text{-}380}$ (AF-P18615) coloured according to pLDDT. W345 is shown as sticks. **(B)** Predicted structure of CBC-NELF-E shown is Figure 1I coloured according to pLDDT. **(C)** Predicted aligned error (PAE) plot for the Alphafold CBC-NELF-E model**.** **(D)** Superposition of the Alphafold-modelled NELF-E residues binding into a groove between CBP80 and CBP20, compared to the experimentally determined X-ray structure of the CBC-NELF-E complex (PDB-code: 5OOB) shown in magenta. The Alphafold model is coloured according to pLDDT. **(E)** Sequence alignment of NELF-E proteins. Only the sequence of residues 344-380 interacting with CBC is shown. Identical residues are in brown boxes. Blue squares indicate residues involved in the

interaction with CBC. **(F)** ITC measurement of the interaction affinity between CBC and NELF-E$^{244-380}$ in the presence of m$^{7}$GTP. **(G)** ITC measurement of the interaction affinity between CBC and NELF-E$^{244-380}$ (W581E) in the presence of m$^{7}$GTP. **(H)** Superdex 200 gel filtration elution profiles of WT and W581E mutation-containing NELF-E$^{244-380}$. The elution profiles are similar, indicating that the W345E mutation does not significantly affect the overall behaviour of this NELF-E fragment. **(I, J)** SDS-PAGE analysis of fractions 1-5 corresponding to the peak of the Superdex 200 gel filtration elution profiles of WT and W345E mutant of NELF-E shown in **H**.

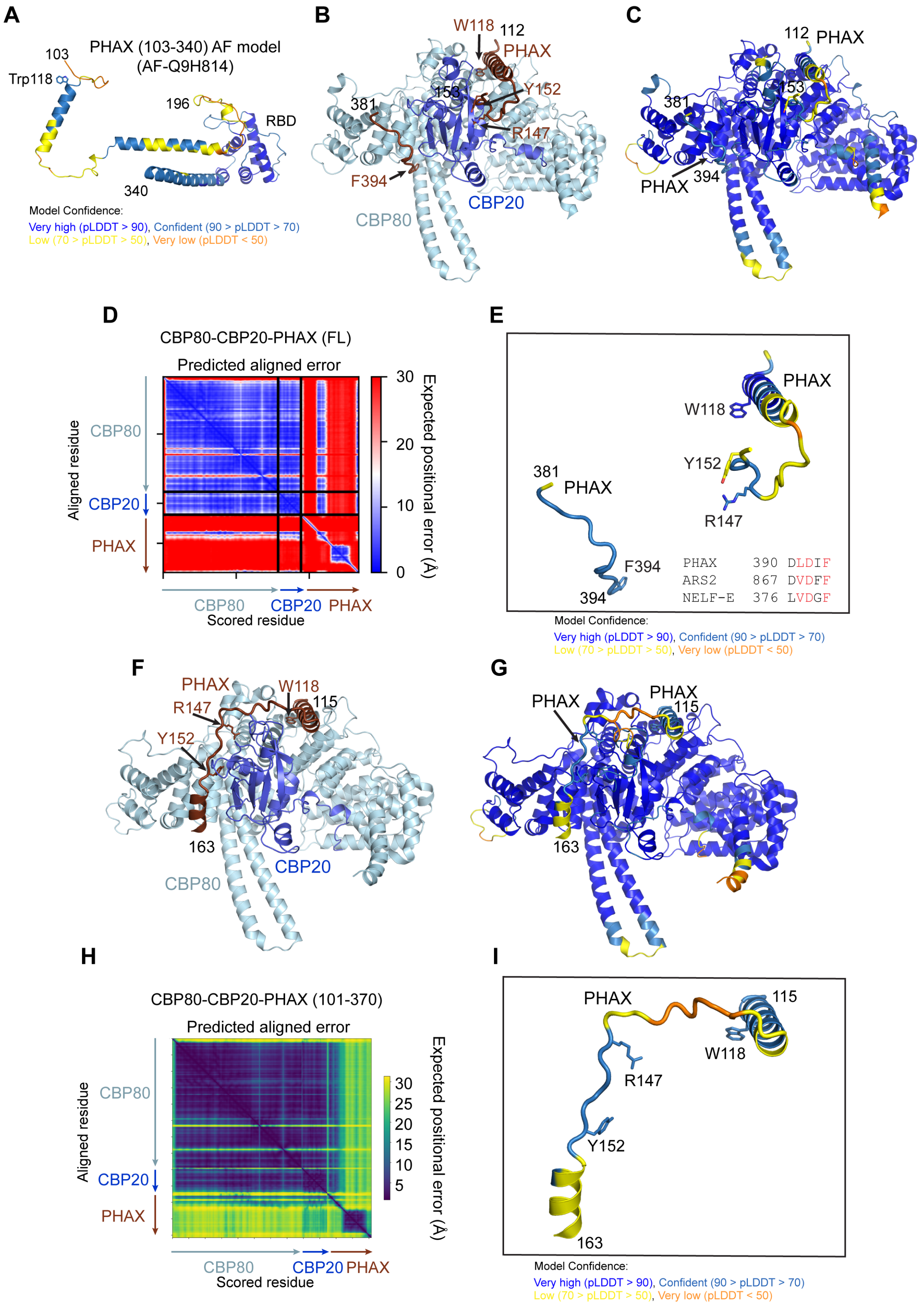


**Figure S5. Alphafold prediction of the CBC-PHAX complex structure.**

**(A)** Alphafold database model of PHAX$^{103\text{-}340}$ (AF-Q9H814) coloured according to pLDDT. W118 is shown as sticks. **(B)** Alphafold-multimer prediction of the structure of the CBC-PHAX complex coloured by chain with PHAX in dark brick. Only residues 112-153 and 381-394 of PHAX are shown. W118, R147, Y152 and F394 of PHAX are shown as sticks. Residues 1-17 of CBP80 predicted with low confidence are not shown. **(C)** Predicted structure of CBC-PHAX coloured according to pLDDT. **(D)** Predicted aligned error (PAE) plot for the AlphaFold CBC-PHAX model shown in **B-C**. **(E)** Predicted structure of PHAX residues 112-153 and 381-394 when bound to CBC, coloured according to pLDDT. Sequence of the PHAX C-terminal motif is compared to the corresponding sequences in ARS2 and NELF-E. **(F)** Alphafold-multimer prediction of the structure of CBP80, CBP20 and $PHAX^{101-370}$. Only residues 115-163 of PHAX are shown. W118, R147 and Y152 of PHAX are shown as sticks. Residues 1-13 of CBP80 predicted with low confidence are not shown. **(G)** Predicted structure of CBC-$PHAX^{101-370}$ coloured according to pLDDT. **(H)** PAE plot for the AlphaFold CBC-PHAX model shown in **F-G**. **(I)** Predicted structure of PHAX residues 115-163 when bound to CBC, coloured according to pLDDT.

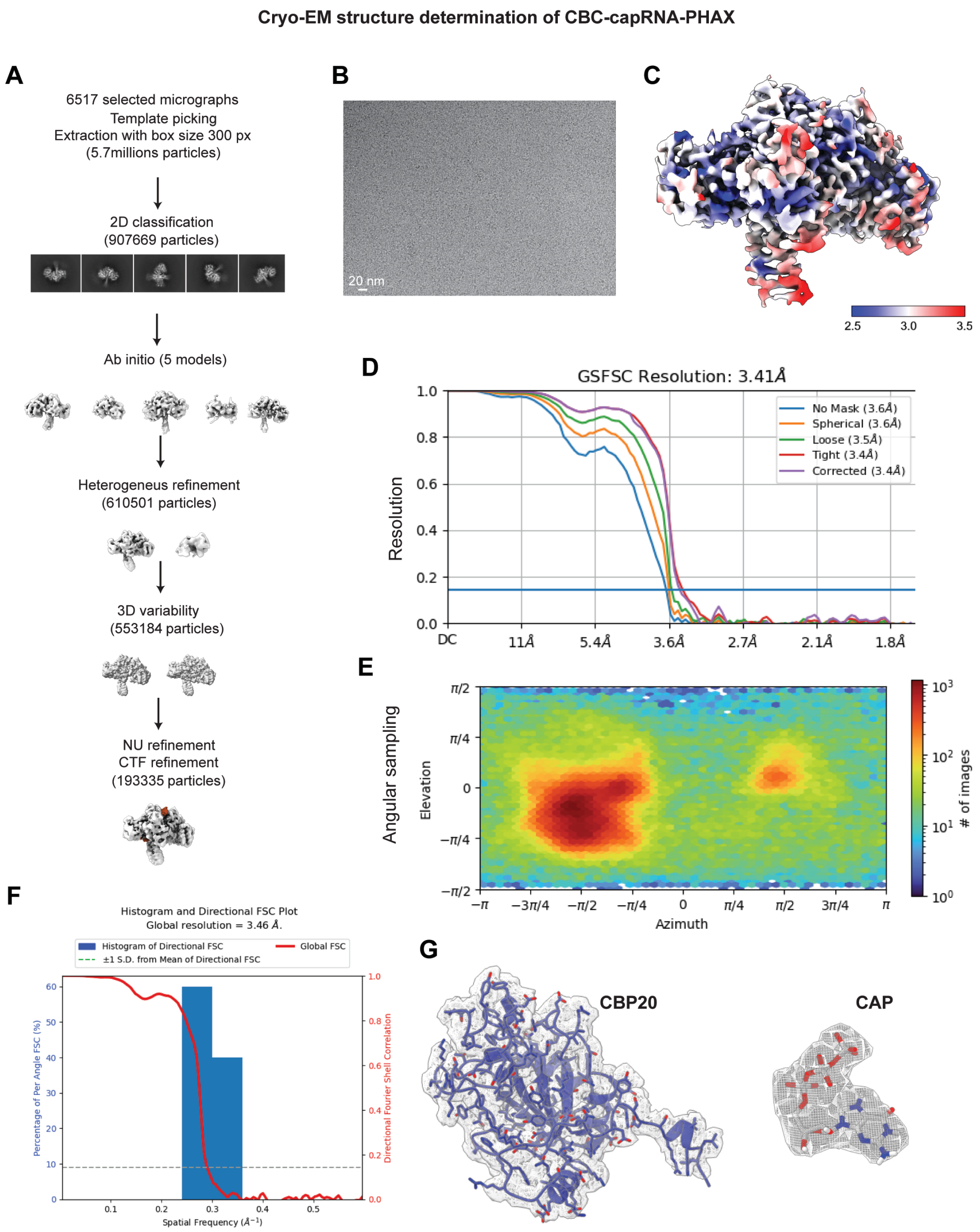


**Figure S6. Cryo-EM analysis of the CBC-capRNA-PHAX complex.**
**(A)** Summary of the cryo-EM processing workflow used to obtain the final map, as described in the Methods section. The cryo-EM maps are unsharpened and the location of the PHAX protein is shown in dark brick. **(B)** Example of an aligned micrograph. **(C)** Final cryo-EM density filtered based on the resolution, sharpened with B-factor -100Å$^2$ and colored according to local resolution. **(D)** Map resolution according to the FSC threshold of 0.143. **(E)** Angular sampling distribution scheme. **(F)** Map resolution according to the 3D FSC. **(G)** cryo-EM density corresponding to CBP20 and m$^7$GpppG.

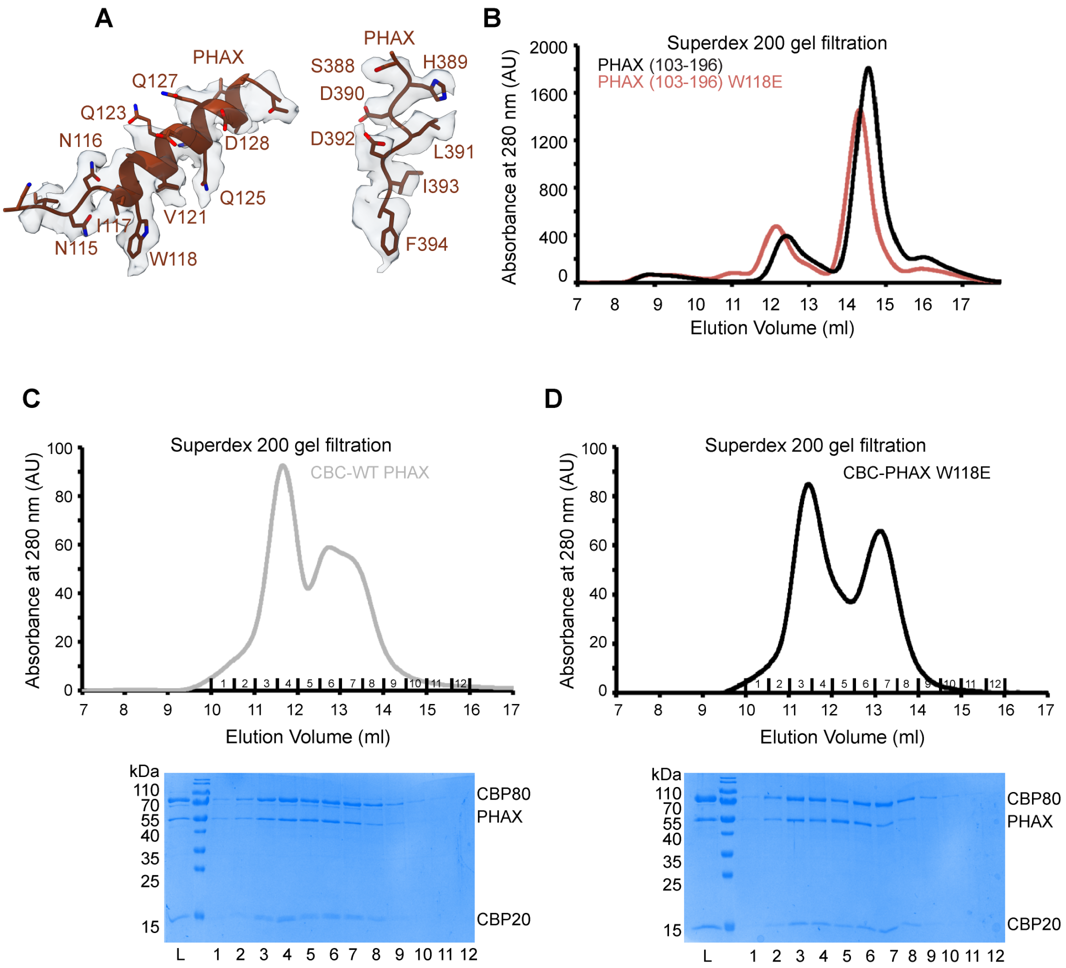


**Figure S7. Characterization of the interaction between CBC and PHAX.**
**(A)** Cryo-EM density for W118-containing helix and the C-terminal peptide of PHAX within the CBC-capRNA-PHAX complex**. (B)** Superdex 200 gel filtration elution profiles of WT and W118E mutation-containing PHAX$^{103\text{-}196}$. The elution profiles are similar, indicating that the W118E mutation does not significantly affect the overall behaviour of this PHAX fragment. **(C)** Superdex 200 gel filtration elution profile of CBC mixed with WT FL His-PHAX. The two proteins co-elute in the first peak. The second peak corresponds to the excess of CBC. Fractions 1-12 were analyzed by SDS-PAGE (lower panel). **(D)** Superdex 200 gel filtration elution profile of CBC mixed with FL His-PHAX (W118E). The two proteins co-elute in the first peak. The second peak corresponds to the excess of CBC. Fractions 1-12 were analyzed by SDS-PAGE (lower panel).

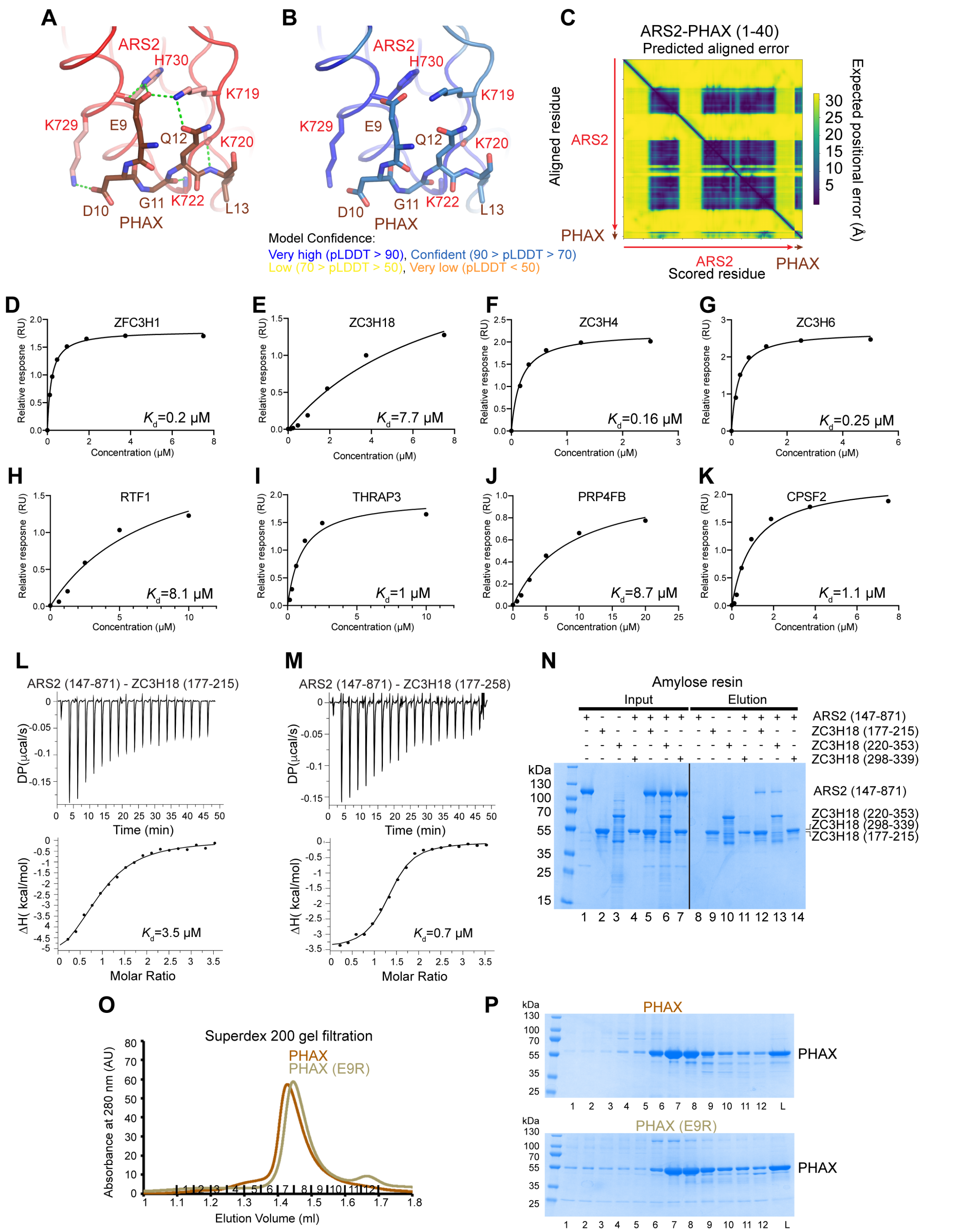

**Figure S8. Characterization of the interaction between ARS2 and RNA effectors.**
**(A)** Details of the interaction between the ARS2 effector domain and the EDGQL motif of PHAX (residues 9-13, in dark brick) as modelled by Alphafold-multimer. The model was predicted for full-length ARS2 and PHAX$^{1-40}$. The model with full-length PHAX revealed the same contacts. **(B)** Details of the Alphafold-multimer modelled interaction between the ARS2 effector domain and the EDGQ motif of PHAX complex coloured according to pLDDT. **(C)** Predicted aligned error (PAE) plot for the Alphafold ARS2-PHAX model shown in **A-B.** **(D-K)** BLI measurement of the binding affinity

between His-ARS2$^{147\text{-}871}$ and the ARM motif containing peptides. $K_D$ are calculated using the integrated Octet RED96e (ForteBio) software with a 1:1 kinetic model and drawn using Graph Pad Prism5. Measurements were performed in duplicate for each peptide. **(L)** ITC measurement of the interaction affinity between ARS2$^{147\text{-}871}$ and His-MBP-ZC3H18$^{177\text{-}215}$. **(M)** ITC measurement of the interaction affinity between ARS2$^{147\text{-}871}$ and His-MBP-ZC3H18$^{177\text{-}258}$. **(N)** MBP pull-down experiments of His-MBP-ZC3H18 constructs with ARS2$^{147\text{-}871}$. All proteins were first purified by affinity chromatography. Proteins were mixed as indicated above the lanes. A total of 4% of the input (lanes 1–7) and 40% of the eluates (lanes 8–14) were analysed on 4-20% gradient SDS-PAGE gels stained with Commassie brilliant blue. ARS2$^{147\text{-}871}$ is retained by ZC3H18$^{117\text{-}215}$ and ZC3H18$^{220\text{-}353}$ (lanes 12,13), but not by ZC3H18$^{298\text{-}339}$ (lane 14). **(O)** Superdex 200 gel filtration elution profiles of WT and E9R mutation-containing PHAX$^{103\text{-}196}$. The elution profiles are similar, indicating that the E9R mutation does not significantly affect the overall behaviour of this PHAX fragment. **(P)** SDS-PAGE analysis of fractions 1-12 corresponding to the peak of the Superdex 200 gel filtration elution profiles of WT and E9E mutant of PHAX shown in **O**.

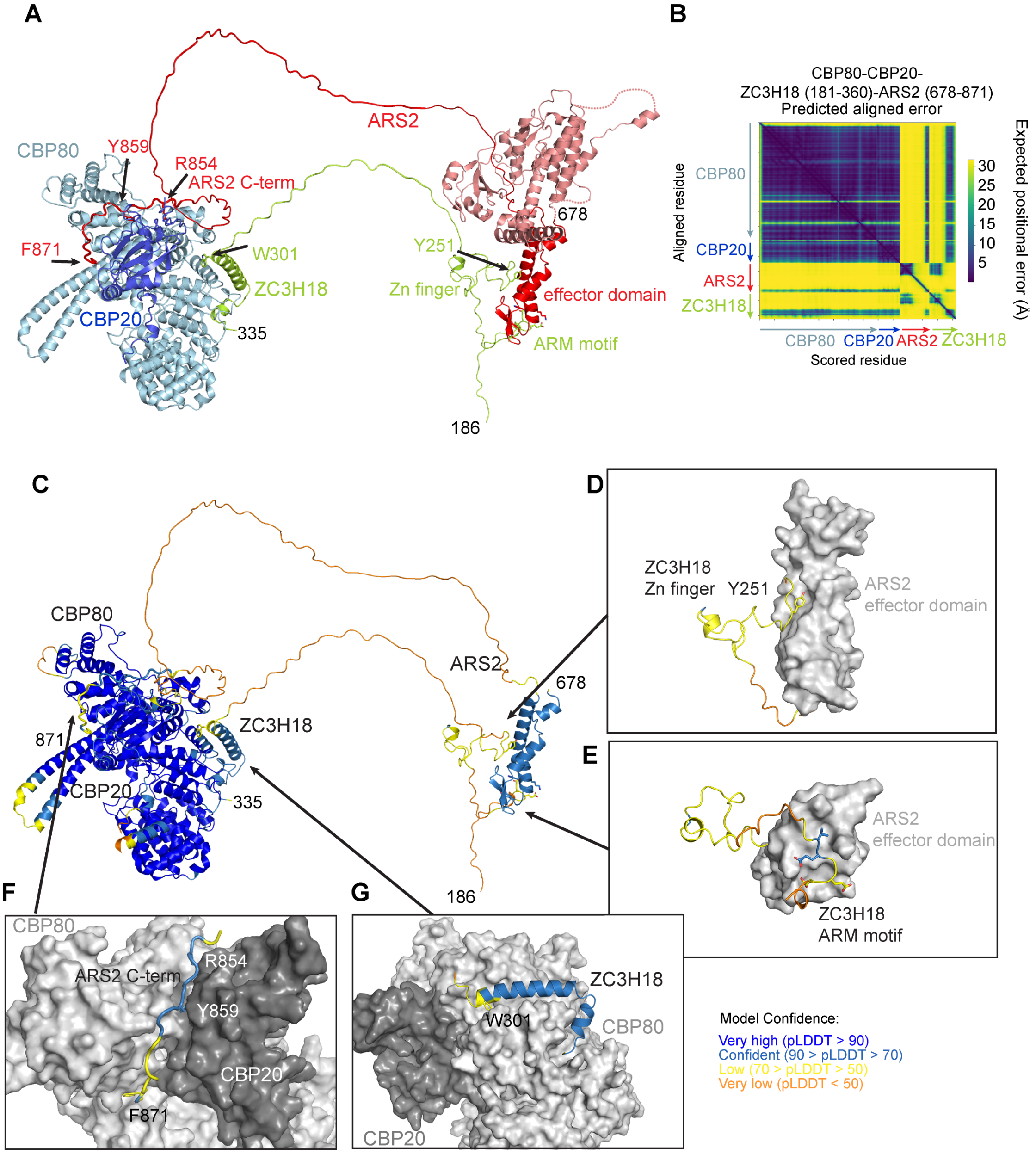


**Figure S9. Alphafold prediction of the CBC-ARS2-ZC3H18 complex structure.**
**(A)** Alphafold-multimer predicted structure of the CBC-ARS2-ZC3H18 complex coloured by chain with ARS2 in red and ZC3H18 in lemon. The model was predicted for CBP80, CBP20, ARS2$^{678\text{-}871}$ and ZC3H18$^{181\text{-}360}$. Only residues 186-335 of ZC3H18 are shown. Residues 1-18 of CBP80 predicted with low confidence are not shown. Key ZC3H18 and ARS2 residues involved in the interactions with their partners are shown as sticks. A crystal structure of the full-length ARS2 is superimposed on the predicted structure of the effector domain of ARS2 (PDB code: 6F7P, shown in pink). **(B)** Predicted aligned error (PAE) plot for the Alphafold CBC-ARS2-ZC3H18 model shown in **A**. **(C)** Predicted structure of the CBC-ARS2-ZC3H18 complex coloured according to pLDDT. **(D)** Details of the interaction between the ZC3H18 peptide 248-KGNYSL (colored by pLDDT) and the shaft of the ARS2 effector domain. **(E)** Details of the interaction between the ZC3H18 ARM motif (colored by pLDDT) and the ARS2 effector domain **(F)** Details of the interaction between the C-terminus of ARS2 (colored by pLDDT) and the groove between CBP80 and CBP20. **(G)** Details of the interaction between the ZC3H18 segment 297-334 (colored by pLDDT) and CBP80.

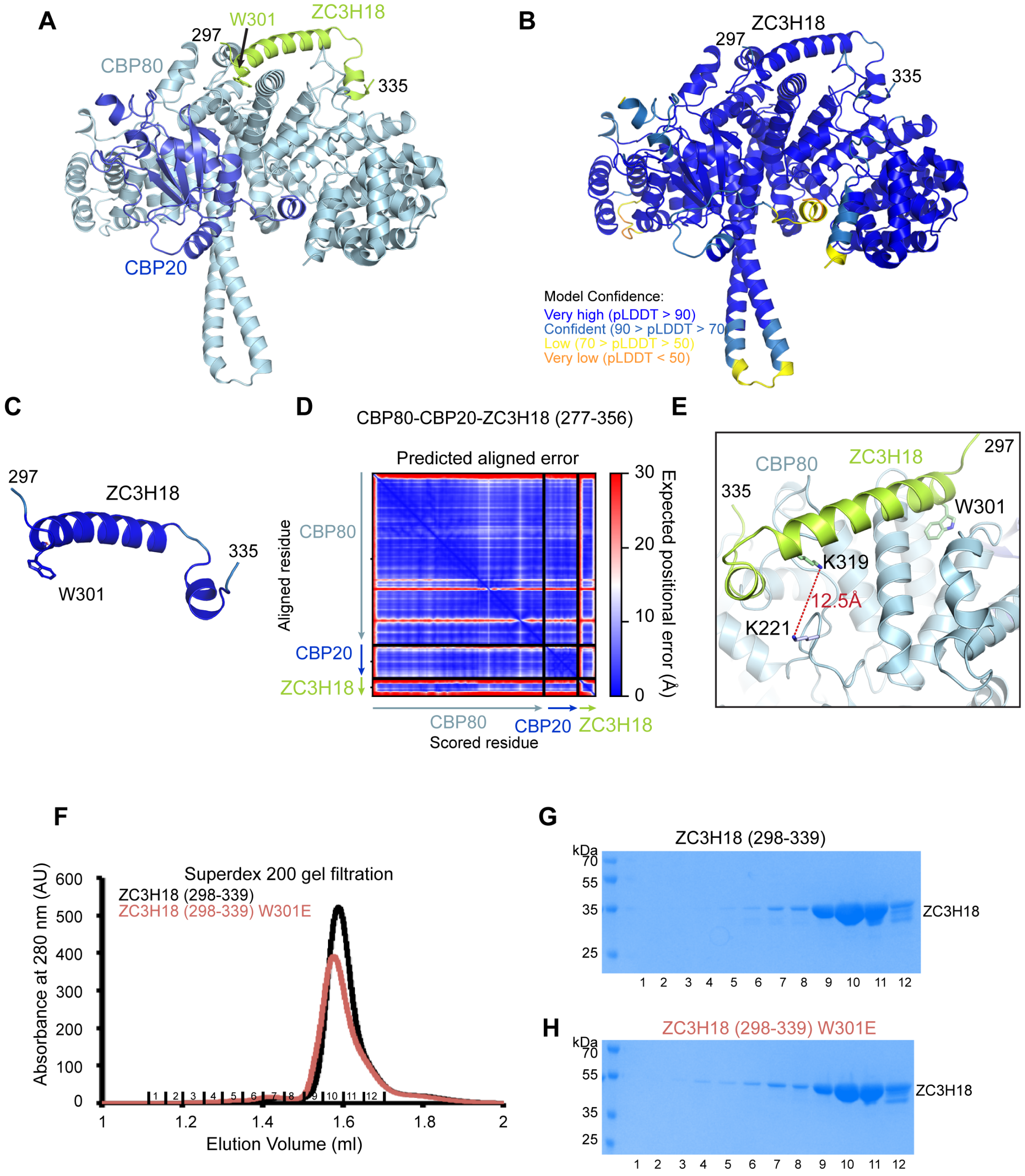


**Figure S10. Alphafold prediction of the CBC-ZC3H18 complex structure.**

**(A)** Alphafold-multimer predicted structure of the CBC-ZC3H18 complex shown in Figure 4B, coloured by chain with ZC3H18 in lemon. The model was predicted for CBP80, CBP20 and ZC3H18$^{277\text{-}356}$. Only residues 297-335 of ZC3H18 are shown. Residues 1-21 of CBP80 predicted with low confidence are not shown. **(B)** Predicted structure of the CBC-ZC3H18 complex coloured according to pLDDT. **(C)** Predicted structure of ZC3H18 residues 297-335 when bound to CBC coloured according to pLDDT. **(D)** Predicted aligned error (PAE) plot for the Alphafold CBC-ZC3H18 model shown in **A-B** and Figure 4B**. (E)** K319 of the predicted CBC-interacting region of ZC3H18 was shown to cross-link with K221 of CBP80 (Townsend et al., 2020). These two residues are predicted to be 12.5Å apart, which corresponds to the crosslinking distance of BS3 used in that study. **(F)** Superdex 200 gel filtration elution profiles of WT and W301E mutation-containing ZC3H18$^{298\text{-}339}$. The elution profiles are similar, indicating that the W301E mutation does not significantly affect the overall behaviour of ZC3H18$^{298\text{-}339}$. **(G, H)** SDS-PAGE analysis of fractions 1-12 corresponding to the peak of the Superdex 200 gel filtration elution profiles of WT and W301E mutant of ZC3H18$^{298\text{-}339}$shown in **F**.

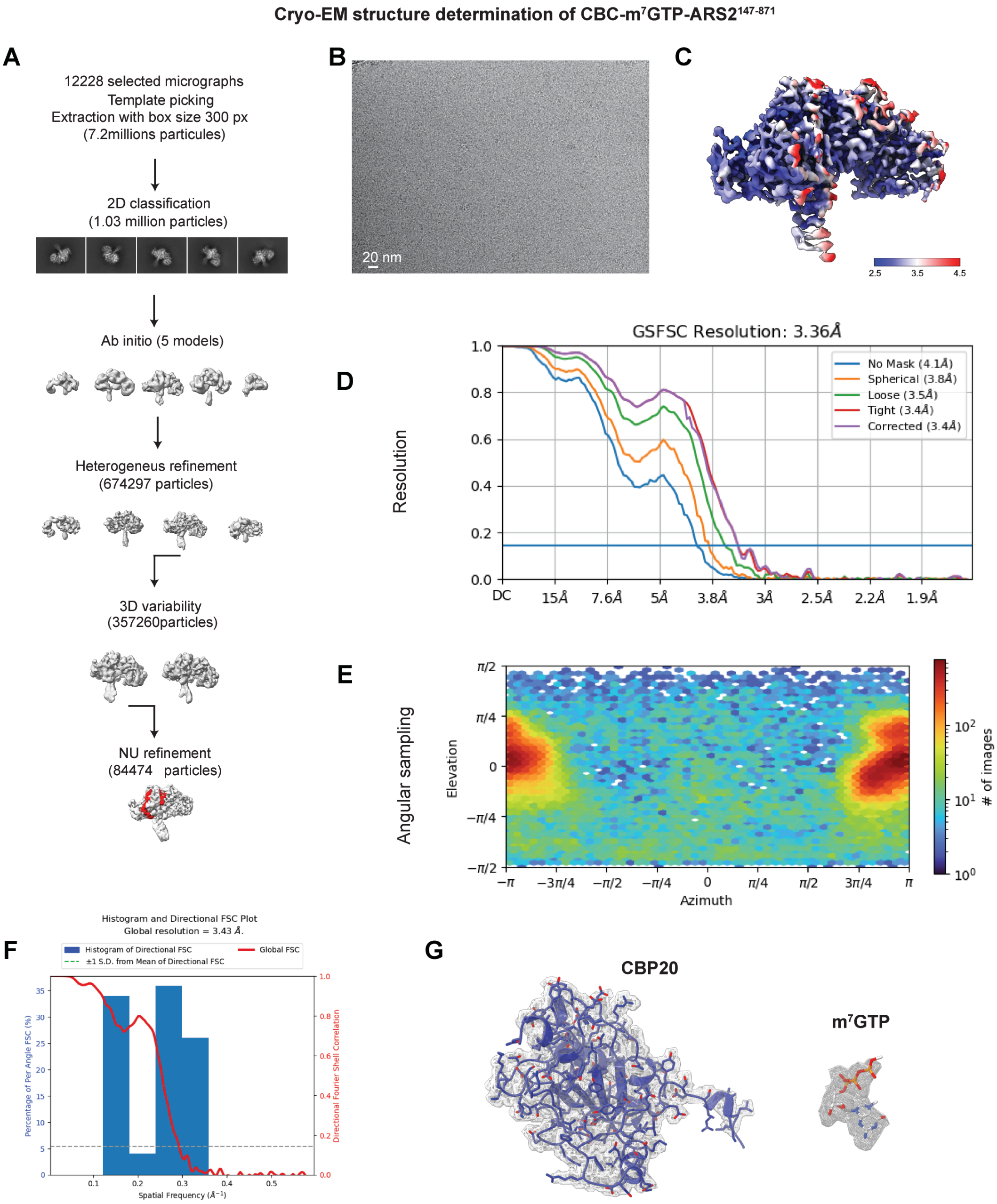


**Figure S11. Cryo-EM analysis of the CBC-m$^7$GTP-ARS2$^{147\text{-}871}$ complex.**

**(A)** Summary of the cryo-EM processing workflow used to obtain the final map, as described in the Methods section. The cryo-EM maps are unsharpened and the location of the ARS2 protein is shown in red. **(B)** Example of an aligned micrograph. **(C)** Final cryo-EM density filtered based on the resolution, sharpened with B-factor -100Å$^2$ and colored according to local resolution. **(D)** Map resolution according to the FSC threshold of 0.143. **(E)** Angular sampling distribution scheme. **(F)** Map resolution according to the 3D FSC. **(G)** Cryo-EM density corresponding to CBP20 and m$^7$GTP.

### Table S1. Cryo-EM data collection, refinement and validation statistics

| | CBC-m7GpppG-NCBP3$^{560-620}$ (PDB-####, EMD-####) | CBC-m7GTP-ARS2$^{147-871}$ (PDB-####, EMD-####) | CBC-capRNA-PHAX (PDB-####, EMD-####) |
|---|---|---|---|
| Short name | CBC-NCBP3 | CBC-ARS2 | CBC-PHAX |
| **Data collection and processing** | | | |
| Date of data collection | 02-04-2021 | 12-05-23 | 07-04-23 |
| Microscope (ESRF) | Krios K2, EF | Krios K3, EF | Krios K3, EF |
| Magnification | 165,000 x | 105,000 x | 105,000x |
| Voltage (kV) | 300 | 300 | 300 |
| Electron exposure (e–/Å$^2$) | 40 | 40 | 40 |
| Defocus range (µm) | -0.7 and -2,5 | -0.7 and -2,5 | -0.7 and -2,5 |
| Pixel size (Å/pix) | 0.87 | 0.84 | 0.84 |
| SuperRes pixel size (Å/pix) | n/a | n/a | n/a |
| Symmetry imposed | - | - | - |
| Initial particle images (no.) | 2773604 | 8095459 | 6400950 |
| Final particle images (no.) | 152563 | 84474 | 193335 |
| Global map resolution (Å) | 3.19 | 3.30 | 3.30 |
| 3DFSC resolution (Å) FSC threshold 0.143 | 3.54 | 3.43 | 3.46 |
| Map resolution range (Å) (25$^{th}$ to 75$^{th}$ percentile) | 2.8-5.8 | 2.8-5.8 | 2.8-5.8 |
| **Refinement and Validation** | | | |
| **PDB file** | | | |
| Map sharpening *B* factor (Å$^2$) | -100 | -100 | -100 |
| Model composition | | | |
| Non-hydrogen atom | 7601 | 7554 | 7609 |
| Protein residues | 923 | 920 | 927 |
| Waters | 0 | 0 | 0 |
| Ligands | 1 (m7GpppG) | 1 (m7GTP) | 1 (m$^7$Gppp-AAUCUAUAAUAG) |
| *B* factor (Å$^2$) | 59.3 | 142.4 | 137.3 |
| R.m.s. deviations | | | |
| Bond lengths (Å) | 0.004 | 0.006 | 0.002 |
| Bond angles (°) | 0.652 | 0.715 | 0.489 |
| Validation | | | |
| MolProbity score | 1.75 | 1.92 | 1.39 |
| Clashscore | 6.96 | 16.53 | 4.51 |
| Poor rotamers (%) | 2.88 | 0.12 | 0.12 |
| Ramachandran plot | | | |
| Favored (%) | 97.92 | 96.81 | 97.05 |
| Allowed (%) | 2.08 | 2.86 | 2.73 |
| Disallowed (%) | 0.00 | 0.33 | 0.22 |
| Model vs Data | | | |
| CC (mask) | 0.82 | 0.68 | 0.83 |
| CC (box) | 0.86 | 0.82 | 0.86 |
| CC (peaks) | 0.78 | 0.59 | 0.74 |
| CC (volume) | 0.81 | 0.67 | 0.82 |
| Mean CC for ligands | 0.77 | 0.60 | 0.71 |

### Table S2: Cell lines used or generated during this study

| Name | Source |
|---|---|
| Mouse ES-E14TG2A *OsTIR1-HA Zc3h18-3F-mAID* | (Polàk et al., 2023) |
| Mouse ES-E14TG2A *OsTIR1-HA Zc3h18-3F-mAID MYC-Zc3h18$^{WT}$* | This study |
| Mouse ES-E14TG2A *OsTIR1-HA Zc3h18-3F-mAID MYC-Zc3h18$^{W297E}$* | This study |
| Mouse ES-E14TG2A *OsTIR1-HA Zc3h18-3F-mAID MYC-Zc3h18$^{ARMmut}$* | This study |

### Table S3: Plasmids used or generated during this study

| Name | Source |
|---|---|
| pCR8[mZC3H18] | This study |
| pB[MYC-ZC3H18] BSD | This study |
| pB[MYC-ZC3H18$^{W297E}$] BSD | This study |
| pB[MYC-ZC3H18$^{D188A, E190A, D193A, E195A, E203A, E205A}$] BSD | This study |

### Table S4: qPCR primers used during this study

| Name | Sequence |
|---|---|
| GAPDH_F | TTGATGGCAACAATCTCCAC |
| GAPDH_R | CGTCCCGTAGACAAAATGGT |
| proDDX56_F | CCCTGACCCACAGAGTGACG |
| proDDX56_R | CCACAGAACCCTAATTCCTTTGCG |
| proRPL27a_F | CGTCGGAGTGCACTGTTCTT |
| proRPL27a_R | GAAGTCTTGCCGATGCTCTG |
| proPURB_F | GACGCTCCCGGTTTCAGAGG |
| proPURB_R | GGAGGTGATTGGCTCCACTG |
| proFIGNL1_F | CTTGGCTTCCCGTTCACTGC |
| proFIGNL1_R | GTGCTCTCTACACTCCTGAGC |
| OCT4_F | CAGCAGATCACTCACATCGCCA |
| OCT4_R | GCCTCATACTCTTCTCGTTGGG |

### Table S5: Antibodies used during this study

| Target | Host | Source |
|---|---|---|
| FLAG (M2) | Mouse | Sigma-Aldrich (#F1804) |
| MYC | Rabbit | Cell Signalling (#2278) |
| ARS2 | Rabbit | GeneTex (#GTX119872) |
| NCBP1 | Rabbit | Abcam (#ab42389) |
| ACTIN | Mouse | Sigma-Aldrich (#A2228) |
| RPLP0 | Rabbit | Abcam (#ab192866) |

### References

Townsend, C., Leelaram, M.N., Agafonov, D.E., Dybkov, O., Will, C.L., Bertram, K., Urlaub, H., Kastner, B., Stark, H., and Luhrmann, R. (2020). Mechanism of protein-guided folding of the active site U2/U6 RNA during spliceosome activation. Science *370*.
